# Cerebral Small Vessel Disease Burden is Associated with Decreased Abundance of Gut Barnesiella intestinihominis Bacterium in the Framingham Heart Study

**DOI:** 10.1101/2022.09.27.509283

**Authors:** Bernard Fongang, Claudia L. Satizabal, Tiffany F. Kautz, Yannick W. Ngouongo, Jazmyn A. SherraeMuhammad, Erin Vasquez, Julia Mathews, Monica Goss, Amy R. Saklad, Jayandra Himali, Alexa Beiser, Jose E. Cavazos, Michael C. Mahaney, Gladys Maestre, Charles DeCarli, Eric L. Shipp, Ramachandran S. Vasan, Sudha Seshadri

## Abstract

A bidirectional communication exists between the brain and the gut, in which the gut microbiota influences cognitive function and vice-versa. Gut dysbiosis has been linked to several diseases, including Alzheimer’s disease and related dementias (ADRD). However, the relationship between gut dysbiosis and markers of cerebral small vessel disease (cSVD), a major contributor to ADRD, is unknown. In this cross-sectional study, we examined the connection between the gut microbiome, cognitive, and neuroimaging markers of cSVD in the Framingham Heart Study (FHS). Markers of cSVD included white matter hyperintensities (WMH), peak width of skeletonized mean diffusivity (PSMD), and executive function (EF), estimated as the difference between the trail-making tests B and A. We included 972 FHS participants with MRI scans, neurocognitive measures, and stool samples and quantified the gut microbiota composition using 16S rRNA sequencing. We used multivariable association and differential abundance analyses adjusting for age, sex, BMI, and education level to estimate the association between gut microbiota and WMH, PSMD, and EF measures. Our results suggest an increased abundance of *Pseudobutyrivibrio* and *Ruminococcus* genera was associated with lower WMH and PSMD (p-values < 0.001), as well as better executive function (p-values < 0.01). In addition, in both differential and multivariable analyses, we found that the gram-negative bacterium *Barnesiella intestinihominis* was strongly associated with markers indicating a higher cSVD burden. Finally, functional analyses using *PICRUSt* implicated various KEGG pathways, including microbial quorum sensing, AMP/GMP-activated protein kinase, phenylpyruvate, and β-hydroxybutyrate production previously associated with cognitive performance and dementia. Our study provides important insights into the association between the gut microbiome and cSVD, but further studies are needed to replicate the findings.

## Introduction

With the increase in life expectancy, age-related diseases are anticipated to cause an enormous burden on health care and social systems. Cerebral small vessel disease (cSVD) contributes to cognitive impairment and dementia in the elderly, with characteristic early deficits in information processing and executive function (EF).^1–6^

Several markers derived from magnetic resonance imaging (MRI) have been established as essential tools for the diagnosis and research on cSVD, including the peak width of skeletonized mean diffusivity (PSMD) and white matter hyperintensity volumes (WMH)^1^,^6–8^. PSMD is a fully automated diffusion tensor imaging (DTI) marker based on the skeletonization of white matter tracts and histograms analysis of mean diffusivity (MD)^9^,^10^. PSMD is highly sensitive to vascular-related white matter lesions and is associated with processing speed in patients with cSVD.^11^ WHM (total and periventricular) are focal and multifocal confluent lesions of white matter injury that reflect axonal loss due to chronic ischemia caused by cSVD^5^,^11^,^12^.Studies have shown that extensive WMH burden is associated with an increased risk of incident stroke and dementia. The initial stage of cSVD is characterized by a decline in EF, commonly observed as a deficit in problem-solving, attention, and decreased information processing speed. Combined with MRI measures of brain structure and neuropsychological assessment, measuring executive function allows for both a structural and functional approach to understanding the etiology of cSVD^13–18^.

A gut microflora imbalance has been linked to several neurological diseases, including Parkinson’s disease, Alzheimer’s disease, and stroke.^19–25^ Studies have shown that the gut microbiome modulates the host brain function through the bidirectional *microbiota-gut-brain* axis via various signaling pathways, including the endocrine, enteric nervous systems, immune systems, and tryptophan metabolism ^19^,^20^,^26–40^. Moreover, it is suggested that the gut microbiota may play a role in vascular cognitive impairment and dementia, as well as related risk factors, such as cSVD. For example, the administration of *Clostridium butyricum*, a butyrate-producing gram-positive bacterium, to vascular dementia mouse models has been shown to attenuate cognitive dysfunction and protect against cerebral ischemia/reperfusion injury.^41^ Gut dysbiosis has also been associated with cSVD, where patients with Enterotype I were more likely to have cognitive decline and high total cSVD scores.^42^ Others have also shown that gut-derived Metabolite Phenylacetylglutamine levels are associated with WMH burden in patients with acute ischemic stroke^43^.

Although studies have reported an association between the gut microbiome and cSVD, whether this association can be extended to structural (i.e., WMH and PSMD) MRI markers of cSVD and cognitive performance (i.e., EF) is unknown.^42^ In this study, we used stool, MRI, and neurocognitive data from the Framingham Heart Study to assess the cross-sectional association between the gut microbiome and markers of cSVD. We found that several phyla, genera, and species were associated with individual cSVD markers. However, only one bacterium, the gram-negative species *Barnesiella intestinihominis*, was negatively associated with all markers of cSVD with a consistent direction of association. In addition, our differential abundance analysis indicated that a higher burden of cSVD and lower EF were associated with decreased abundance of *Barnesiella intestinihominis*. Finally, we did not find statistical differences in gut microbiome diversity in FHS participants stratified by cSVD burden groups.

## Material and Methods

### Sample Description

The FHS is an ongoing population-based, longitudinal cohort initiated in 1948, recruiting 5,209 participants from Framingham (MA, USA). The FHS’s initial aim was the prospective investigation of the risk factors associated with CVD and has since expanded to other diseases. In 1971, 5,214 children of the Original Cohort (the Offspring Cohort) and their spouses were enrolled and underwent similar examinations as the Original Cohort every four years. The children of the Offspring participants and their spouses have been followed since 2002 as the 3^rd^ Generation cohort, and participants have been examined on several occasions. The current project was limited to participants from the New Offspring Spouse, the 3^rd^ Generation, and the OMNI 2 cohorts who attended the third examination (2016-2019) and provided their stool specimens. All participants provided written informed consent for blood drawn, MRI testing, and cognition assessments at each examination. The IRB of Boston University School of Medicine approved the FHS protocols and participant consent forms.

#### MRI acquisition and processing

Participants were imaged on a 1.5 or 3T scanner. We used 3D-T1, fluid-attenuated inversion recovery (FLAIR), and Diffusion Tensor Imaging (DTI) sequences to derive the neuroimaging markers. Image analyses were performed at the Imaging of Dementia and Aging (IDeA) laboratory at UC Davis, using established pipelines.^44^ All images were analyzed by operators blinded to all participant characteristics, including cognitive performance on neuropsychological testing. Peak width of skeletonized mean diffusivity (PSMD) is a novel and fully automated MRI biomarker, which has shown clinical relevance in SVD, an essential contributor to VCI.^45^,^46^ It is based on skeletonization and histogram analysis of diffusion imaging DTI data. PSMD scores have also been shown to be strongly correlated with processing speed.^47^ PSMD is measured as the difference between the 5^th^ and 95^th^ percentiles of the distribution of the voxel MD value across the skeleton of the brain white matter. PSMD and WMH measures were expressed as the percentage of total intracranial volume to correct for head size and were further log-transformed due to skewness before analyses.

#### Executive Function

Clinical neuropsychologists and trained research assistants administered validated neuropsychological tests. We derived the difference between the Trail Making Test B and Trail Making Test A (Trails B-A) as a measure of EF.^48^,^49^ Values of Trails B-A were natural log-transformed to normalize its distribution before analyses and further inversed such that higher EF scores indicate poor performance. Details of the neuropsychological protocol can be found elsewhere.^48^,^49^

### Microbiome

#### Sample Handling and DNA Extraction

Stool samples were collected in 100% ethanol as previously described and stored at −80°C.^50^,^51^ For DNA and RNA co-extraction, the QIAamp 96 PowerFecal Qiacube HT Kit (Qiagen Cat No./ID: 51531) was paired with the Allprep DNA/RNA 96 Kit (Qiagen Cat No./ID: 80311), and IRS solution (Qiagen Cat No./ID: 26000-50-2) for a custom protocol. For initial lysis, 50 - 200 mg of stool per sample were frozen into individual wells of the PowerBead plate containing 0.1 mm glass beads (Cat No./ID: 27500-4-EP-BP) on a dry ice block. Next, 650 μl of 55°C heated PW1 buffer and 25 μL of freshly-prepared 1M DTT were added directly to each sample well before lysis by beating on a TissueLyzer II at 20 Hz for a total of 10 minutes (in two 5-minute intervals with plate rotation in between). Next, samples were pelleted by centrifugation for 6 minutes at 4,500 x g, and supernatants were transferred to a new S block (supplied in PowerFecal Kit), combined with 150 μl of IRS solution, and vortexed briefly before a one-minute incubation. Next, sealed samples were centrifuged again for 6 minutes at 4,500 x g, and up to 450 μl of supernatant was transferred to a new S block, combined with 600 μl of Buffer C4 (PowerFecal Kit), mixed by pipetting ten times and incubated for 1 minute. Next, samples were transferred into an AllPrep 96 DNA plate on clean S blocks and centrifuged for 3 minutes at 4,500 x g. This centrifugation step was repeated until the entire sample had been centrifuged. Finally, the AllPrep 96 DNA plate was stored at 4°C until after DNA extraction.

The Allprep 96 DNA plate was removed from 4°C and placed on top of a 2mL waste block for DNA extraction. First, 500 μl AW1 buffer was added to the DNA plate and sealed before centrifuging for 4 minutes at 4,500 x g. The waste block was emptied after each wash step. Next, 500 μl AW2 buffer was added to the DNA plate, sealed with AirPore tape, and centrifuged for 10 minutes at 4,500 x g. Next, the Allprep 96 DNA plate was placed on the elution plate, and 100 μl of 70°C heated EB Buffer was added to each sample column and incubated for 5 minutes. The DNA plate was sealed and then centrifuged for 4 minutes at 4,500 x g to elute the DNA, which was stored at −20°C. All incubation and centrifugation steps were performed at room temperature.

#### 16S rRNA Gene Sequencing

16S rRNA gene libraries targeting the V4 region were prepared by first using qPCR to normalize template concentrations and determine the optimal cycle number. Library construction was performed in quadruplicate with the primers 515F (5’-AATGATACGGCGACCACCGAGATCTACACTATGGTAATTGTGTGCCAGCMGCCGCGGTA A-3’) and unique reverse barcode primers from the Golay primer set (Caporaso, 2011 and 2012). After amplification, sample replicates were pooled and cleaned via the Agencourt AMPure XP-PCR purification system. Prior to final pooling, purified libraries were normalized via qPCR in two 25 μL reactions, 2x iQ SYBR SUPERMix (Bio-Rad, REF: 1708880) with Read 1 (5’-TATGGTAATT GT GTGYCAGCMGCCGCGGTAA-3’), Read 2 (5’-AGTCAGTCAG CC GGACTACNVGGGTWTCTAAT-3’) primers. Pools were quantified by Qubit (Life Technologies, Inc.) and sequenced on an Illumina MiSeq with 2 x 150 bp reads using custom index 5’-ATTAGAWACCCBDGTAGTCC GG CTGACTGACT-3’ and custom Read 1 and Read 2 primers mentioned above.

### Microbiome Data Analysis

We used QIIME2 ^52^ for downstream processing of microbiome sequencing data. Briefly, forward and reverse reads were truncated to preserve minimum *Phred* quality scores of 28 in 75% of reads. Next, high-quality sequencing reads were clustered into operational taxonomic units (OTUs) at a 97% dissimilarity threshold. Then, the clustered OTUs were classified against the Greengenes database to assign taxonomy to the sequences. Lastly, we constructed phylogenetic trees using the MAFFT algorithm in QIIME2. To avoid deviation caused by the effects of different sequencing depths and low representatives, we removed OTUs with fewer than four reads in less than 10% of samples. We also constructed a table of relative abundance counts without rarefaction for multivariate analyses. Finally, we exported the OTUs, taxonomy, phylogeny, and metadata tables to the R platform for further research. Finally, the cross-associations between MRI measures and gut microbial composition were assessed using multivariate linear models, considering MRI measures as continual variables, and differential analyses in which MRI measures were divided into burden groups.

### Microbiome multivariable linear regression

Multivariable association analyses were conducted using the R package MaAsLin2^53^ using relative abundance counts without rarefaction and the different diversity indexes. We used the following model

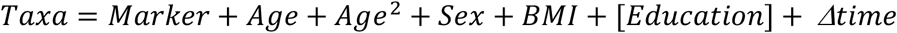

Where *Taxa* is relative abundance counts, *Marker* is the log-transformed cSVD marker (WMH, PSMD, EF) measure, and *Δtime* is the time difference between the stool sample collection and the MRI scan. In addition, we considered the age at which MRI scans were completed, and only EF measures were adjusted for the participant’s education level. Finally, MaAsLin2 analysis was done using the following settings: minimum abundance (relative) cutoff at 10^-3^, the minimum percent of samples for which a feature is detected at minimum abundance was set a 10%, and we selected the negative binomial as the regression method.

### Microbiome differential abundance analysis

We used the following procedure to stratify participants into cSVD burden groups according to WMH and PSMD markers, and a similar strategy was used for cognitive function (EF): (**i**) cSVD markers were log-transformed to ensure normality; (**ii**) for each marker, we created two groups of interest dichotomized by the upper quintile (20% of participants, *“**High burden”*** group) and the bottom four quintiles (80% of participants, *“**Lower burden”*** group) of the residual. The differential analysis of microbiome composition analyses was conducted by comparing differences in abundance and alpha-diversity (Observed, Chao1, Shannon, and Simpson index) using a repeated-measures analysis of variance (ANOVA). Beta-diversity was estimated using principal coordinated analysis (PCoA) with Bray-Curtis, weighted and unweighted UniFrac distances. PCoA is an unsupervised dimensionality reduction method that can be used to visualize group separations of compositional data. Finally, we used permutational analysis of variance (PERMANOVA) implemented in the vegan package to test for differences in community composition among the *“**Lower burden”*** and ***“High burden”*** groups, followed by pairwise tests between groups. The microbiome *uniqueness*, which reflects how dissimilar individuals are from their neighbors in the population, was computed as in Wilmanski et al.^54^ Statistically significant differences were reported after adjusting for multiple testing using the Benjamini-Hochberg procedure. We conducted the differential abundance analysis using MaAsLin2, with relative abundance counts, adjusting for relevant covariates (age, age^2^, sex, BMI, education, and *Δtime).^53^* For taxonomy features significantly different between groups, we performed functional profiling of the bacterial metagenome using the Phylogenetic Investigation of Communities by Reconstruction of Unobserved States (*PICRUSt*).

## Results

### 1 Study population

A total of 1,406 participants successfully returned their stool collection kits. Of these, 9 samples failed the quality control process after DNA extraction and sequencing, and 417 participants did not have MRI or neurocognitive measures. Therefore, the present study was limited to the 980 participants who were administered MRI scans in a previous examination or during the visit at which stool samples were requested. In addition, we excluded participants with dementia (3) and incident stroke (5) and, thus,retained 972 participants for further analyses. Participants’ characteristics are presented in **Table 1**. The study flowchart is provided as supplementary material (**Supplementary File SF1, Figure S1).** In addition, a correlation plot depicting the relationship between cSVD markers and the different covariates is provided in **Supplementary File SF1, Figure S2**.

**Table 1:** Descriptive table of demographics variables.

| Characteristic | N = 972<sup>1</sup> |
| --- | --- |
| AGE | 54 (47, 60) |
| SEX, F | 540 (56%) |
| BMI | 27.0 (24.2, 31.0) |
| EDUC |  |
| No High School Degree | 3 (0.3%) |
| High School Degree | 78 (8.0%) |
| Some college | 203 (21%) |
| College Graduated | 573 (59%) |
| WMH |  |
| Range | -3.35 (-4.20, -2.49) |
| Lower Burden | 832 (86%) |
| High Burden | 136 (14%) |
| PSMD |  |
| Range ($\times 10^{-3}$) | 0.2 (0.2, 0.3) |
| Lower Burden | 674 (86%) |
| High Burden | 108 (14%) |
| EF |  |
| Range | -1.08 (-1.15, -0.99) |
| Low Burden | 822 (87%) |
| High Burden | 121 (13%) |

#### High cSVD burden is associated with decreased Bacteroidetes abundance

We performed multivariable linear regression analyses to test the associations between the relative abundance of the gut microbiome taxa and cSVD markers (PSMD, WMH, and EF). Statistically significant (adjusted p-values < 0.05) associations are summarized in **Table 2**, and detailed association results are provided in **Supplementary File SF2, Tables 1-3**. At the phylum level, measures of all cSVD markers were negatively correlated with *Bacteroidetes* and positively correlated with *Proteobacteria*, both gram-negative bacteria. Additionally, the grampositive *Firmicutes* was positively correlated with PSMD and EF but negatively correlated with measures of WMH volumes. Finally, the gram-negative *Synergistetes* showed a positive correlation with measures of PSMD and no association with other markers.

**Table 2:**
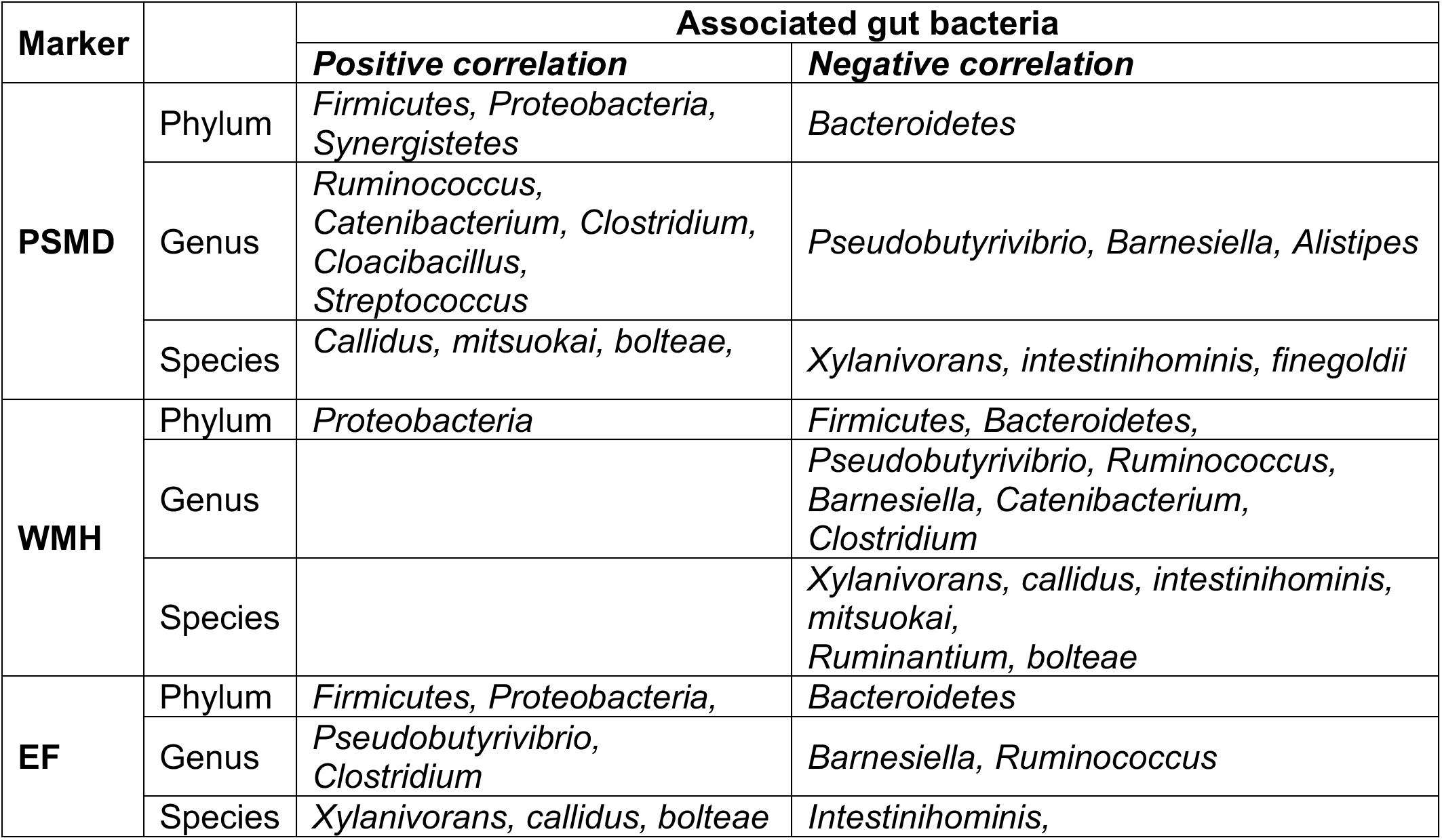
Multivariable association between the gut microbiome and cSVD markers (PSMD, WMH, EF). Statistically significant (BH-adjusted p-value <0.05) bacteria associated with markers of cSVD (PSMD, WMH, EF) were identified through multivariable analysis. The entire table, including p-values and coefficient of association, is provided as supplementary material (**Supplementary File SF2, Tables 1-3**)

*Barnesiella*, a lowly abundant bacterium (<1% of the human gut), was negatively correlated with all measures of cSVD markers at the genus level. In addition, the abundance of the gramnegative *Pseudobutyrivibrio* was positively correlated with measures of EF but negatively correlated with MRI markers PSMD and WMH. *Ruminococcus* was negatively correlated with WMH and EF but positively associated with PSMD. Furthermore, the gram-positive *Clostridium* was positively correlated with EF and PSMD and negatively correlated with WMH volumes. Other genera were positively correlated with PSMD, including *Cloacibacillus* and *Streptococcus*.At the species level, *Xylanivorans*, a butyrate-producing bacterium from the rumen, was negatively correlated with PSMD and WMH and positively correlated with EF. The gram-negative *Intestinihominis*, a species from the *Barnesiella* genus, was negatively correlated with all three measures of cSVD markers. The gram-positive *bolteae* was positively correlated with EF and PSMD and negatively correlated with WMH volumes. Additional species correlated positively (*Callidus, mitsuokai*) and negatively (*finegoldii*) with PSMD. Finally, we also found a negative correlation between WMH volumes and the abundance of *mitsuokai* and *callidus* species.

Altogether, these results suggested an association between cSVD markers’ measures and the gut microbiome’s relative abundance. Therefore, we next performed differential analyses to assess the difference in microbiome composition between FHS participants stratified by cSVD markers burden groups.

#### Differential abundance analysis of the gut microbiome and cSVD markers

Study participants were stratified by burden groups (***Lower*** and ***High burden***),as explained in the methods section. Then, we conducted differential abundance analyses to assess microbial composition differences between groups. The results are summarized in **Table 3,**and the complete list of bacteria differentially abundant between Healthy (Lower burden) and Unhealthy (High burden) groups is provided in **Supplementary File SF2, Tables 4-6.**

**Table 3:** Differential abundance analysis of the gut microbiome and cSVD markers (PSMD, WMH, EF). Statistically significant (BH-adjusted p-value < 0.05) bacteria associated with markers of cSVD (PSMD, WMH, EF) were identified through differential abundance analysis. The whole table, including p-values and coefficient of association, is provided as supplementary material (**Supplementary File SF2, Tables 4-6**)

| Marker |  | Associated gut bacteria |  |
| --- | --- | --- | --- |
|  |  | Increased abundance | Decreased abundance |
| PSMD | Phylum |  | OD1 |
|  | Genus | *cc_115*, *Oscillospira*, *Christensenella* | *Barnesiella*, *Ruminococcus*, *Methanobrevibacter* *Clostridium*, *Victivallis* |
|  | Species | *Spiroforme*, *aldenense*, *guilliermondii*, *ruminantium*, *variabile* | *Intestinihominis*, *lactaris*, *lavalense*, *vadensis* |
| WMH | Phylum | *Proteobacteria* | OD1, *Actinobacteria*, *Bacteroidetes*, *Firmicutes* |
|  | Genus | *Christensenella* | *Ruminococcus*, *Barnesiella*, *Methanobrevibacter*, *Oscillospira*, *cc_115*, *Anaerostinus*, *Clostridium* |
|  | Species | *aldenense* | *Lactaris, intestinihominis, glycerini, ruminantium* |
| EF | Phylum |  | *Firmicutes, OD1* |
|  | Genus | *Collinsella* | *Barnesiella, Methanobrevibacter, Ruminococcus, Alistipes* |
|  | Species | *Aldenense, ruminantium, lavalense, aerofaciens* | *Intestinihominis, lactaris,* |

At the phylum level, the *Candidate Phylum OD1 bacteria (OD1*) had reduced relative abundance (adjusted p-value < 0.05) in the Unhealthy groups compared to Healthy (**Supplementary File SF1 Figures 2-4**) for all three cSVD markers. We also observed a reduced abundance of the gram-positive *Firmicutes* in Unhealthy groups of WMH and EF. Additionally,*Actinobacteria* and *Bacteroidetes* had reduced abundance, and *Proteobacteria* had increased abundance in the Unhealthy WMH group.

**Figure 1:**
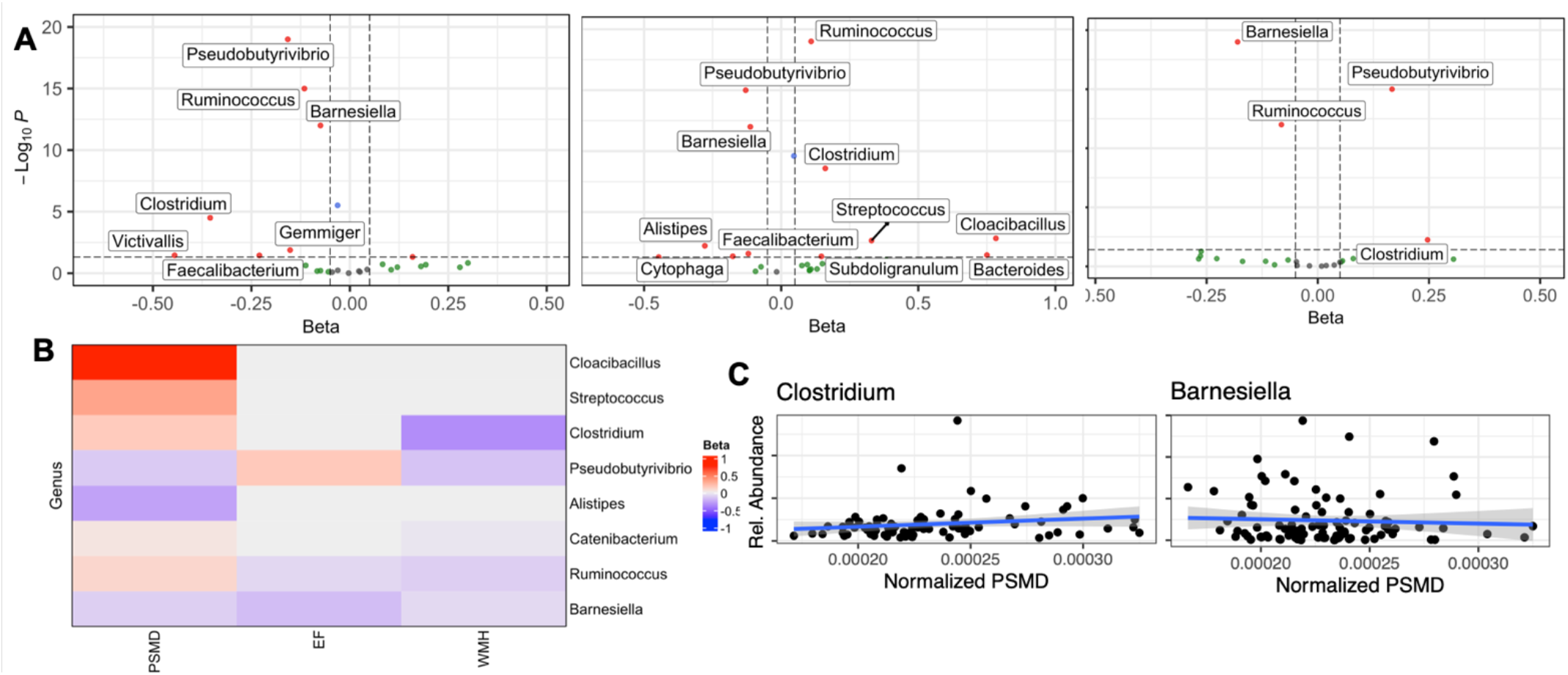
Multivariable association between the gut microbiome and cSVD markers (PSMD, WMH, EF). (A) Volcano plots depicting a significant association (BH-adjusted p-value < 0.05) of several genera and PSMD, WMH, and EF (from left to right). (B) Heatmap of common statistically significant genera associated with cSVD markers. (C) Scatter plots of genera *Clostridium* and *Barnesiella* relative abundance and normalized PSMD measures.

**Figure 2:**
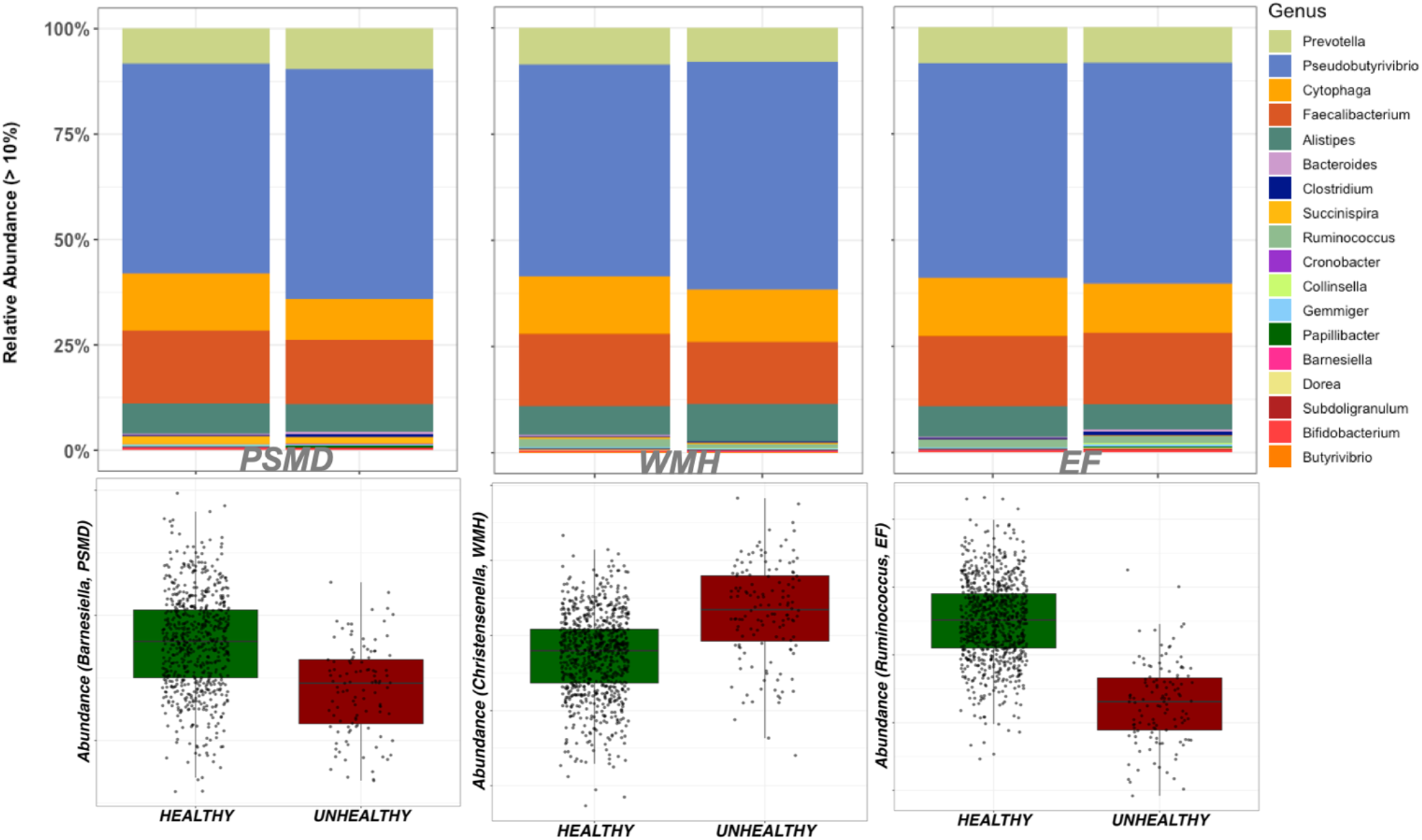
Differential abundance analysis of the gut microbiome and cSVD markers (PSMD, WMH, EF). (top) Stacked plots at 10% minimum abundance highlight differences at the genus level between Healthy and Unhealthy groups of cSVD. (down) relative abundance of *Barnesiella, Christensenella*, and *Ruminococcus* genera in PSMD, WMH, and EF burden groups, respectively (Healthy = Lower burden, Unhealthy = High burden).

At the genus level, the relative abundance of genera *Barnesiella*,*Ruminococcus*, and *Methanobrevibacter* were significantly reduced in the Unhealthy groups of WMH, PSMD, and EF. In addition,*Clostridium*, a gram-positive bacterium that includes several human pathogens, had reduced abundance, while *Christensenella* had increased abundance in Unhealthy PSMD and WHM groups. The genera *cc_115* and *Oscillospira* have increased abundance in PSMD but decreased abundance in WMH Unhealthy groups. Other genera associated with measures of cSVD markers included *Victivallis* (Decreased abundance in PSMD),*Anaerosinus* (decreased abundance in PSMD),*Alistipes* (decreased abundance in EF), and *Collinsella* (Increased abundance in EF).

At the species level,*Intestinihominis* and *lactaris* had decreased abundance in Unhealthy groups, whereas *Aldenense* has increased abundance in Healthy groups of PSMD, WMH, and EF. In addition,*ruminantium* was found to have increased abundance in PSMD and EF but decreased abundance in WMH. Furthermore,*lavalense*, a gram-positive bacterium from the *Clostridium* genius, has increased abundance in EF but decreased in PSMD. Finally, other species associated with cSVD markers in differential abundance analysis included *Aerofaciens* (increased abundance in EF),*vadensis* (decreased abundance in PSMD, and *Spiroforme, guilliermondii*, and *variabile* with increased abundance in PSMD.

The complete list of differentially abundant taxa in cSVD stratified groups is provided in **Supplementary File SF2 Table 4-6;**the box plots of the significant (adjusted p-value < 0.05) taxa are provided in **Supplementary File SF1 Figures 2-4**.

#### *Barnesiella intestinihominis* showed a consistent direction of association in both differential and multivariable association analyses

Multivariable association and differential abundance analyses identified different bacteria associated with markers of cSVD. We hypothesized that bacteria showing a consistent direction of association for both methods are more likely to be accurate. At the genus level,*Barnesiella* was negatively correlated with all cSVD markers and has decreased abundance in Unhealthy individuals, thus supporting the reliability of these associations with cSVD markers. We observed a similar trend at the species level, where the gram-negative *intestinihominis* has decreased abundance in all cSVD markers. Both observations led us to conclude that Unhealthy measures of cSVD (PSMD, WMH, and EF) markers in the FHS are associated with a reduced abundance of *Barnesiella intestinihominis*.

#### cSVD markers do not associate with gut microbiome diversity

##### Biodiversity analysis: α-diversity and β-diversity

We used multivariable and differential analyses to assess the association between cSVD markers and gut microbiome alpha and beta diversity. Measures of α-diversity included observed species, Chao1, Shannon, and Simpson diversity index. Association analyses were adjusted for sex, age, BMI, education, and the time difference between the visits at which data were collected. For differential analysis, we used a one-way analysis of variance (ANOVA) to compute the statistical difference between burden groups (Lower burden, High burden). We did not observe statistical differences between alpha diversity indexes and measures of cSVD markers in multivariable or differential analysis (**Supplementary Figures 17-20**). To estimate the change in the diversity of OTUs between cSVD markers burden groups, we computed the beta diversity and calculated differences using PCoA, as shown in **Supplementary Figures 21-23**. Overall, the different burden groups of all cSVD markers exhibited similar distribution leading to the conclusion that there are very few species differences between samples of FHS participants. Using multivariable analysis, we found that only the Bray-Curtis minimum dissimilarity index (*min_bray*) significantly correlated with WMH (**Supplementary Figure 20**), whereas *min_wnifrac* did not associate with cSVD markers.

#### Predictive functional profiling of microbial communities associated with cSVD

We assessed the potential functional role of the microbial communities associated with cSVD markers using the *PICRUSt* approach described in the Methods. *PICRUSt* predictions were based on KEGG orthologs (*KO*), Enzyme Commission (*EC*) numbers, and MetaCyc metabolic pathways (*MePath*) enrichments, comparing Healthy to Unhealthy measures of cSVD markers. The predicted MePath associated with PSMD included *L-tyrosine, L-methionine, L-phenylalanine*, and *teichoic acid biosynthesis*. In addition, several KO were associated with PSMD, including *adenosine kinase, hemerythrin*, and *glv operon transcriptional regulator* (**Supplementary File 2, Tables 7-9, Supplementary File 1, Figure 24**). Predicted MePath associated with WMH included *taurine* and *adenosine degradation*, significant KOs were related to a *methyltransferase, motility quorum sensor regulator*, and ECs included *Nitrate reductase, hydrogenase, and gluconate 2-dehydrogenase*. Detailed results of the functional profiling of gut bacteria associated with WMH are provided in **Figure 25** of **Supplementary File 1 and Tables 10-12 of Supplementary Figure 2.**For the EF, we found that KOs terms *bacterial CydC and CydD, methyltransferase*, and *type 1 pantothenate kinase*, MePath terms *taxadiene biosynthesis, pyruvate dehydrogenase*, and *reductive TCA cycle-1* were all significantly different between Lower and Higher burden EF groups (**Supplementary File 1, Figure 26**; **Supplementary File 2, Tables 13-15**). Overall, the functional modules associated with the *PICRUSt* predicted metagenome included AMP/GMP-activated kinase, xanthine, homogentisate, tyrosine, phenylpyruvate, and dicarboxylate-hydroxybutyrate. A cross-comparison of the enriched terms between all cSVD markers revealed that common KOs (**Figure 3**) belong to KEGG modules with genes involved in AMP/GMP-activated protein kinase, phenylpyruvate, and β-hydroxybutyrate production.

**Figure 3:**
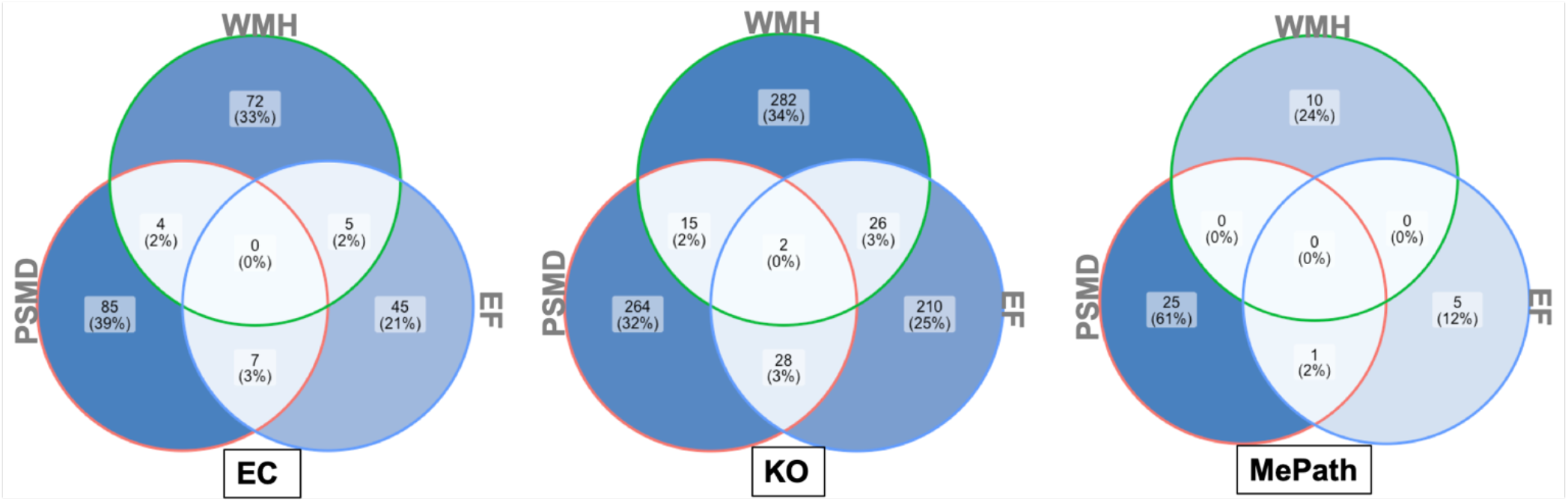
Functional profiling of microbial communities associated with markers of cSVD. *PICRUSt* predictions were based on KEGG Orthologs (KO), Enzyme Commission (EC), and MetaCyc pathways (MePath). The common KOs belong to KEGG modules with genes involved in AMP/GMP-activated protein kinase, phenylpyruvate, and β-hydroxybutyrate production, which have been shown to the associated with cognitive decline and dementia.

## Discussion

In this cross-sectional association study, we used stool samples, MRI (PSMD, WMH), and neurocognitive assessment (Trail making tests A and B) data from 972 middle-aged participants of the FHS to assess the association between the gut microbiome and markers of cSVD. We found that several phyla, genera, and species were associated with individual markers of cSVD, but overall, the gram-negative *Barnesiella intestinihominis* has decreased relative abundance in High burden measures of all cSVD markers.

Our results indicated that measures of PSMD positively correlated with the abundance of *Firmicutes* and *Proteobacteria* but negatively correlated with *Bacteroidetes*. However, in the differential analysis comparing Healthy to Unhealthy participants stratified by PSMD measures, only the *Candidate Phylum OD1 bacteria (OD1*) was found to have decreased abundance in the Unhealthy group. Conversely, elevated WMH measures, predictive of incident stroke, MCI, and dementia, were associated with increased *Proteobacteria* and decreased *Firmicutes* and *Bacteroidetes* in multivariable and differential abundances analyses. Despite the discrepancy in their direction of association, these phyla were previously implicated in cognitive performance.^55^,^56^ Also, the gram-negative *Bacteroidetes*, which excretes pro-inflammatory endotoxin Lipopolysaccharide (LPS), was previously associated with AD in animal and human studies with a contradictory direction of association.^57–61^ Furthermore, the abundance of phylum *Firmicutes, Proteobacteria*, and *Proteobacteria* were also observed in patients with mild cognitive impairment without dementia.^42^,^62^

We also found several genera were associated with markers of cSVD, including *Ruminococcus, Catenibacterium, Clostridium, Cloacibacillus, Alistipes*, cc_115,*Victivallis*, Streptococcus, *Pseudobutyrivibrio*, *Barnesiella*, *Methanobrevibacter*, *Oscillospira*, *Anaerosinus*, and *Collinsella*. Moreover, the genus *Barnesiella has* a consistent direction of association with all cSVD markers. Our results were consistent with previous studies reporting the association between these genera and normal cognition, mild cognitive impairment (MCI), and AD.^56,63,64^ Also, decreased abundance of *Barnesiella* has been previously associated with age-related cognitive decline in mice ^63^. Finally, the relative abundance of *Barnesiella* was shown to correlate with cognitive ability in patients with Parkinson’s disease,^64,65^ and differentially abundant in mice after induced traumatic brain injury.^66^

At the species level, our results indicated an association between cSVD markers and several taxa, including *Callidus, intestinihominis, guilliermondii, Lactaris, ruminantium, Xylanivorans, and bolteae*. Consistent with the observation at the genus level, we found that *intestinihominis* has decreased abundance in the Unhealthy group in all cSVD markers.

Our results suggested that the functional modules associated with *PICRUSt* predicted metagenome included AMP/GMP-activated kinase, xanthine, homogentisate, tyrosine, phenylpyruvate, and dicarboxylate-hydroxybutyrate. These findings align with the known functional roles of the related metabolites in neurodegenerative diseases. Indeed, AMP-activated kinase is a metabolic biosensor with anti-inflammatory activities previously associated with AD, and its activation was proposed to be involved in the elimination of post-synaptic proteins.^67–71^ Similarly, hydroxybutyrate production inhibited inflammasome activation, attenuated AD pathology in mice, and improved cognition in memory-impaired adults.^72,73^

This study has several limitations, and the results should be replicated and mechanistically validated before clinical interpretation. First, the stool samples, MRI scans, and neuropsychological batteries were not collected simultaneously. Although we adjusted for the time difference in our analyses, further studies where all relevant data are collected simultaneously are required. Second, it is widely known that medication and diet influence the gut microbiota and cSVD. In our study, we did not adjust for diet or medication intake as they were unavailable for the studied sample. Further, we may have omitted relevant confounders not accounted for in this analysis, potentially biasing the interpretation of our results. However, these limitations are compensated by the large sample size and the statistical approaches used to assess the link between the gut microbiome and cSVD markers. The statistically significant taxa we reported showed a similar trend in both multivariable association and differential abundance analyses.

## Conclusion

Cerebral small vessel disease is a major cause of cognitive impairment and dementia and is associated with differences in the gut microbiota. In this cross-sectional study, we used data from 972 participants of the Framingham Heart Study to assess whether the gut microbiota is also associated with MRI (PSMD, WMH) and neurocognitive (EF) markers of cSVD. In multivariable and differential analyses, we found the abundance of several phyla, including *Firmicutes, Proteobacteria*, and *Bacteroidetes*, associated with unhealthy measures of cSVD makers. This observation was extended at the genus with *Pseudobutyrivibrio, Barnesiella*, and at the species level with *Intestinihominis, Xylanivorans*, and *lactaris*. Altogether, our results suggest that the gram-negative bacterium *Barnesiella intestinihominis* is strongly associated with unhealthy measures of cSVD markers. In addition, functional analyses indicated that the differentially abundant bacteria gene contents were involved in AMP/GMP-activated kinase, xanthine, homogentisate, tyrosine, phenylpyruvate, and dicarboxylate-hydroxybutyrate processes, all of which were previously associated with cognitive impairment and dementia. Our study provides important insights into the association between the gut microbiome and cSVD, but further studies are needed to replicate the results.

## Author contributions

BF, CS, and SS conceived the study. CS, JH, and AB prepared the data. BF performed all statistical analyses. BF wrote the manuscript. All authors discussed the results, provided feedback during the writing process, and commented on the final manuscript.

## Acknowledgments

The Authors thank Dr. Ramnik Xavier from the Broad Institute of MIT and Harvard, and the Center for Microbiome Informatics and Therapeutics (Massachusetts Institute of Technology, Cambridge, MA, USA) for providing access to the FHS microbiome data.

This study was funded in part by the UT Health San Antonio Center for Biomedical Neuroscience (CBN) and grants from the NIA (AG059421, AG054076, AG049607, AG033090, AG066524, P30 AG066546, 5P30AG059305-03, RF1 AG061729A1, 5U01AG052409-04) and NINDS (NS017950, UF1NS125513, K01NS126489). In addition, Drs. Fongang, Kautz, Seshadri, Satizabal, Maestre, Cavazos, and Himali are partially supported by the South Texas Alzheimer’s Disease Research Center (P30AG066546). Drs. Seshadri and Himali receive support from The Bill and Rebecca Reed Endowment for Precision Therapies and Palliative Care. Dr. Himali is supported by an endowment from the William Castella family as William Castella Distinguished University Chair for Alzheimer’s Disease Research, and Dr. Seshadri by an endowment from the Barker Foundation as the Robert R Barker Distinguished University Professor of Neurology, Psychiatry and Cellular and Integrative Physiology.

## Tables and Figures

### Supplementary File SF1

**Figure S1: Study flowchart**

**Figure S2: cSVD markers and covariates distribution**

**Figure S3: Multivariate association analysis:**volcano plots of PSMD

**Figure S4: Multivariate association analysis:**volcano plots of WMH

**Figure S5: Multivariate association analysis:**volcano plots of EF

**Figure S6-8: Multivariate association analysis:**Scatter plots of PSMD depicting statistically significant associations at the phylum, genus, and species levels

**Figure S9-11: Multivariate association analysis:**Scatter plots of WMH depicting statistically significant associations at the phylum, genus, and species levels

**Figure S12-14: Multivariate association analysis:**Scatter plots of EF depicting statistically significant associations at the phylum, genus, and species levels

**Figure S15-17: Differential abundance analysis:**Differential abundance of the gut microbiome and measures of PSMD, WMH, EF

**Figure S18-20: Alpha diversity measures** (ACE, Chao1, Observed, Shannon, and Simpson indexes) for EF, PSMD, WMH stratified by burden groups

**Figure S21: Correlation between measures of beta-diversity**(min_bray, min_wunifrac), alpha-diversity (ACE, Observed, Chao1, Simpson, and Shannon indexes), and cSVD markers after adjusting for age, age2, sex, BMI, and the time difference between the stool collection and MRI scans

**Figure S22: Principal Coordinate Analysis** depicting the diversity distribution between the different PSMD burden groups

**Figure S23-25: Predicted functional role of the microbial communities associated with PSMD, WMH, EF**

### Supplementary File SF2

**Tables S1-S3**: **Multivariate association analysis:**Statistically significant (adjusted p-values < 0.05) taxa associated with PSMD, WMH, and EF.

**Tables S4-S6**: **Differential abundance analysis:**Statistically significant (adjusted p-values < 0.05) taxa associated with PSMD, WMH, and EF.

**Tables S7-S15**: Functional analysis results with PICRUSt

