## Supplementary File SF1 for "Cerebral Small Vessel Disease Burden is Associated with Decreased Abundance of Gut Barnesiella intestinihominis Bacterium in the Framingham Heart Study"

**Supplementary File SF1: Supplementary Figures**

#### 1. Study Flowchart

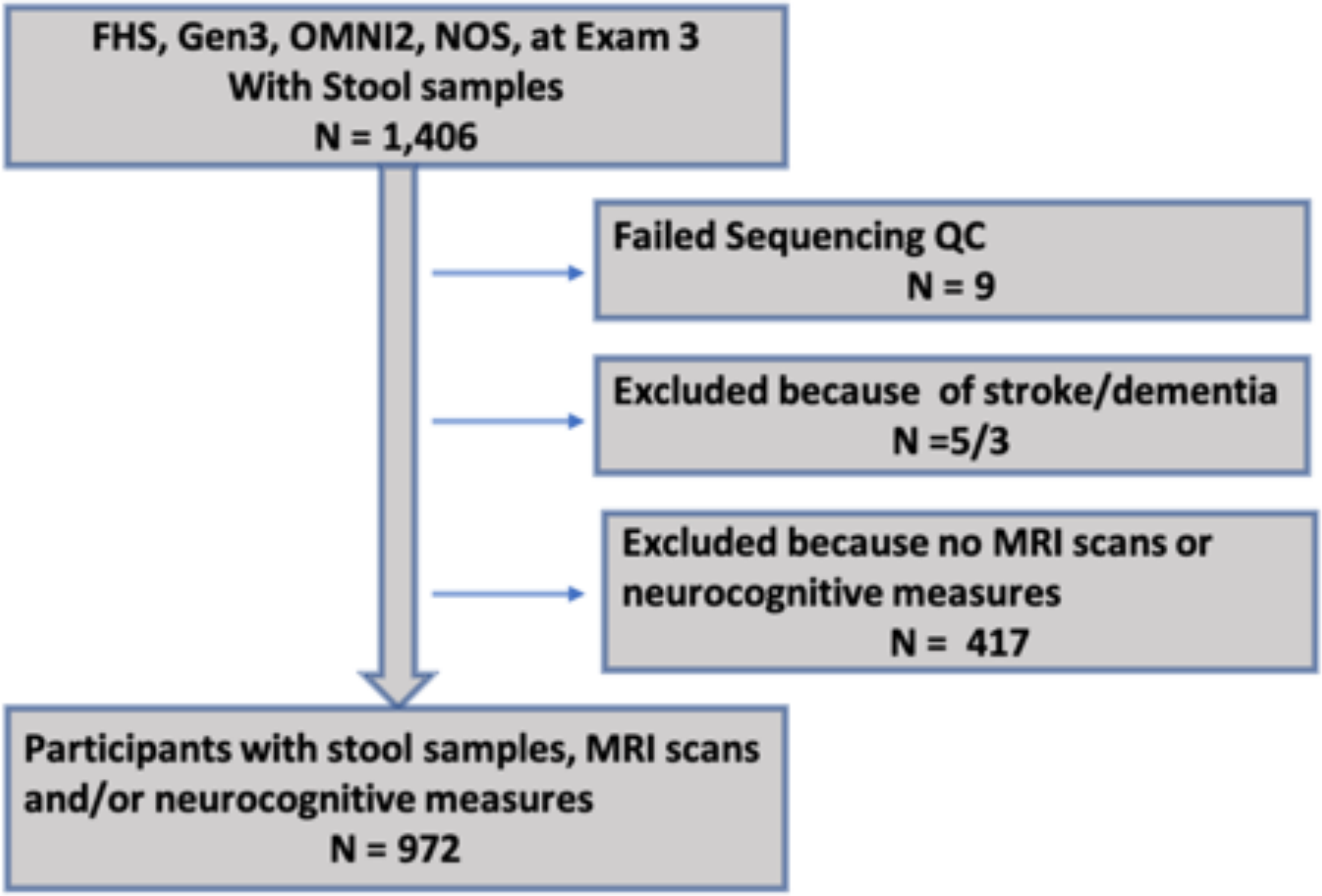

**Figure S1: Study flowchart**

#### 2. cSVD markers and Covariates dependence

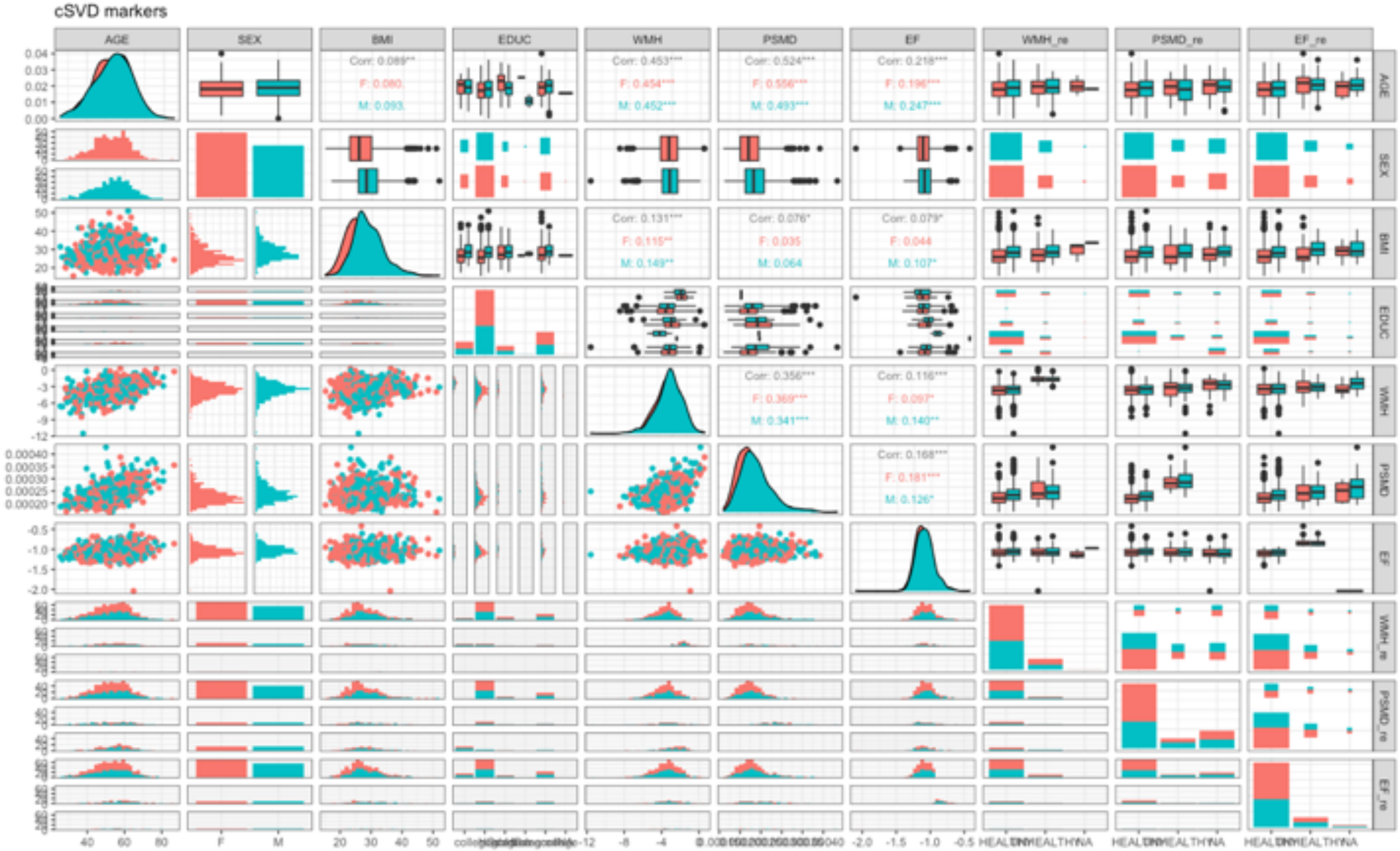

**Figure S2: cSVD markers and covariates distribution.** PSMD_re, WMH_re, and EF_re are stratified measures of PSMD, WMH, and EF, respectively, as described in the text.

#### 2. Multivariable association analysis

##### 2.1 Volcano plots

###### 2.1.1 PSMD volcano plots

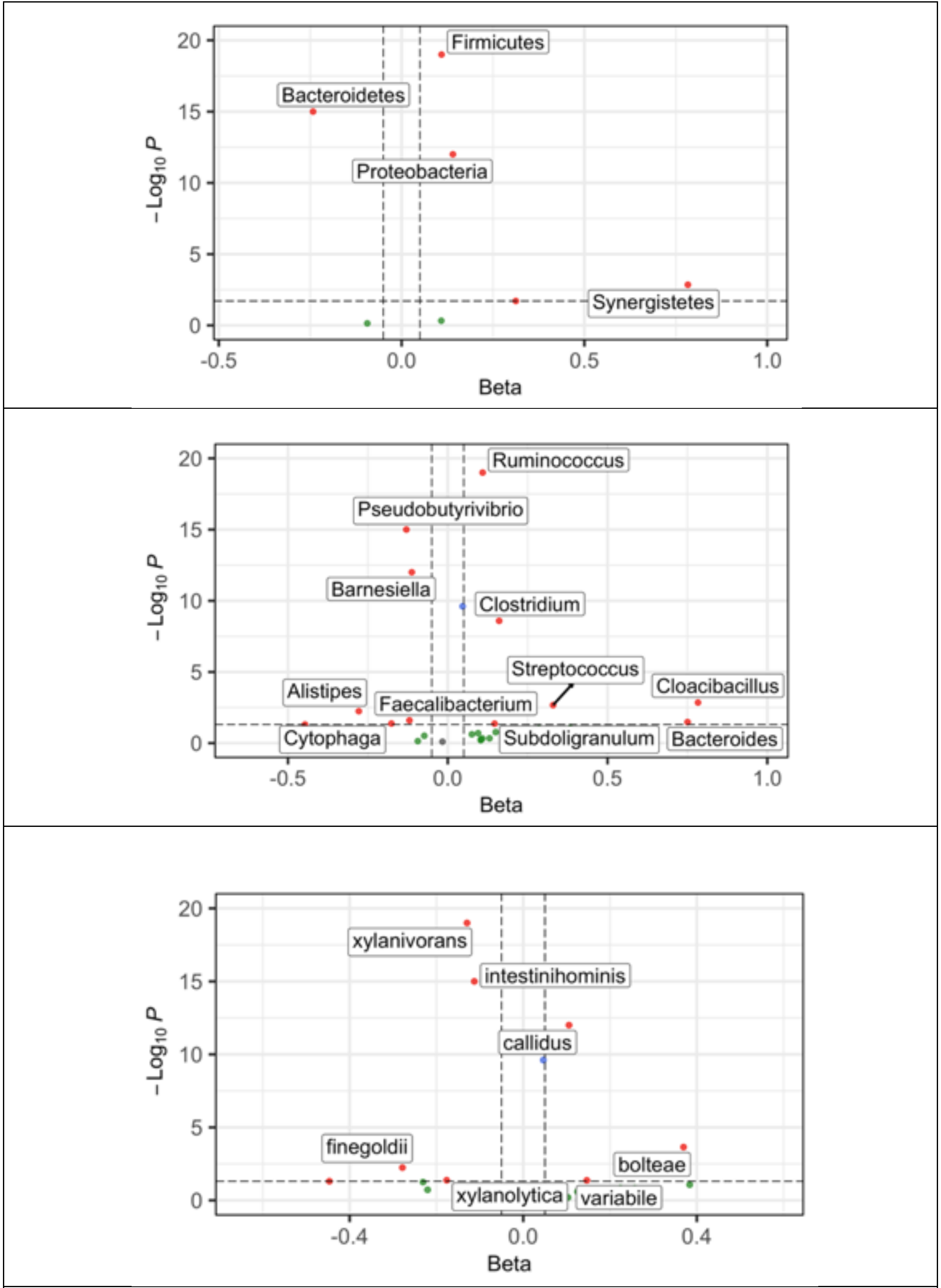

**Figure S3: Multivariate association analysis:** volcano plots of PSMD depicting associations at the phylum, genus, and species levels (top to down)

#### 2.1.2 WMH volcano plots

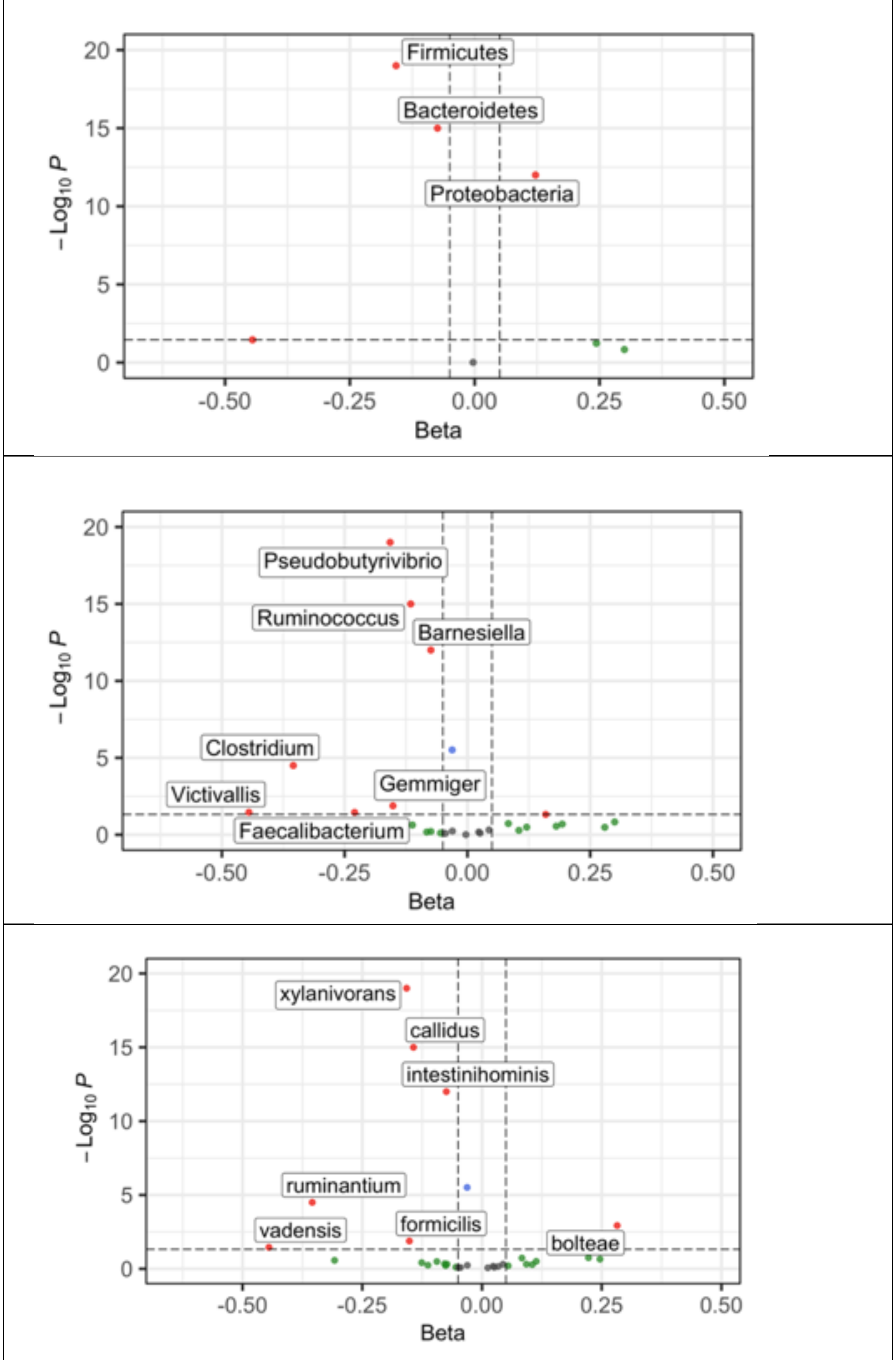

**Figure S4: Multivariate association analysis:** volcano plots of WMH depicting associations at the phylum, genus, and species levels (top to down)

#### 2.1.3 EF volcano plots

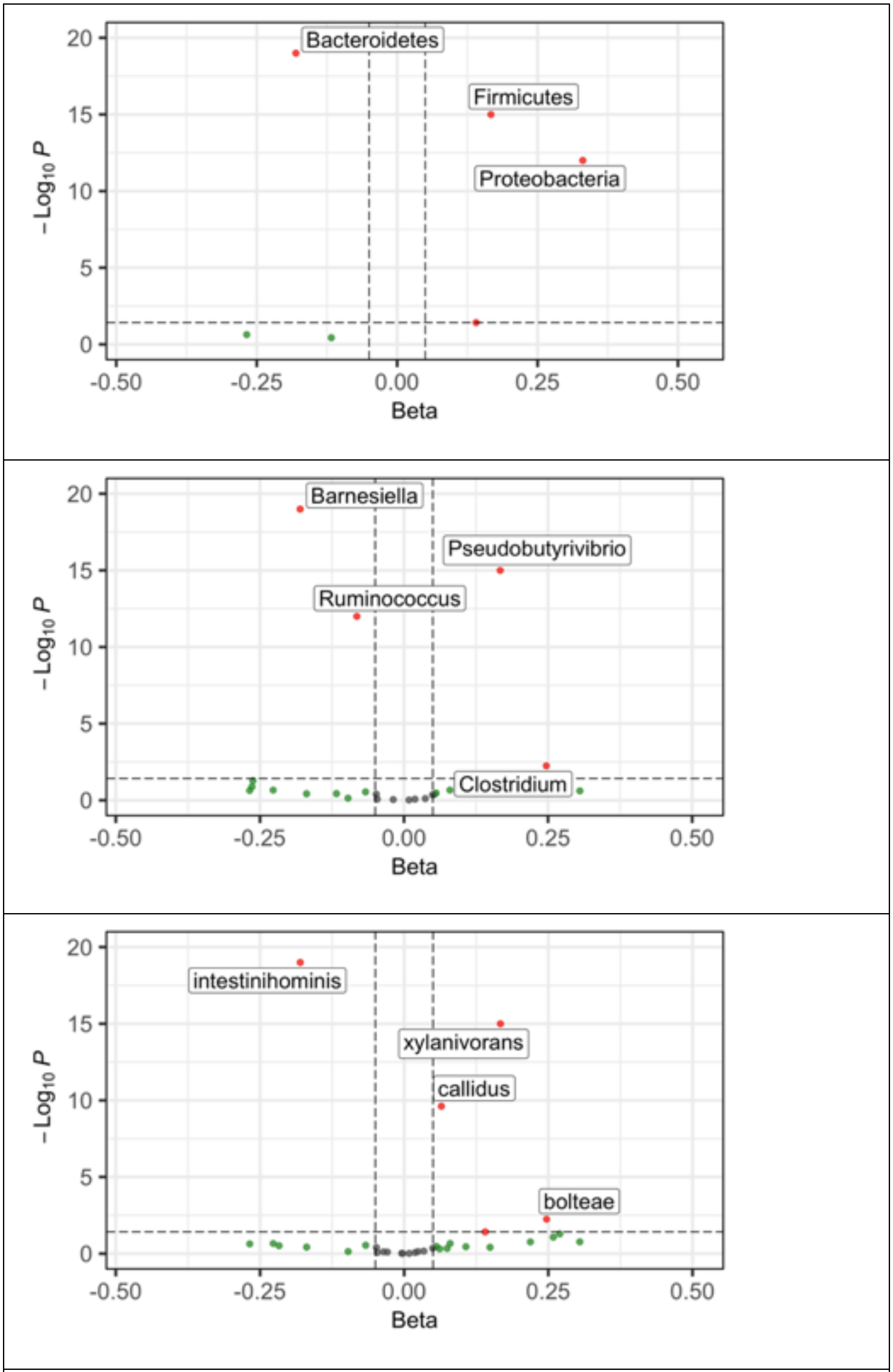

**Figure S5: Multivariate association analysis:** volcano plots of EF depicting associations at the phylum, genus, and species levels (top to down)

#### 2.2 Scatter plots of significant taxa

##### 2.2.1 Scatter plots of significant taxa associated with PSMD

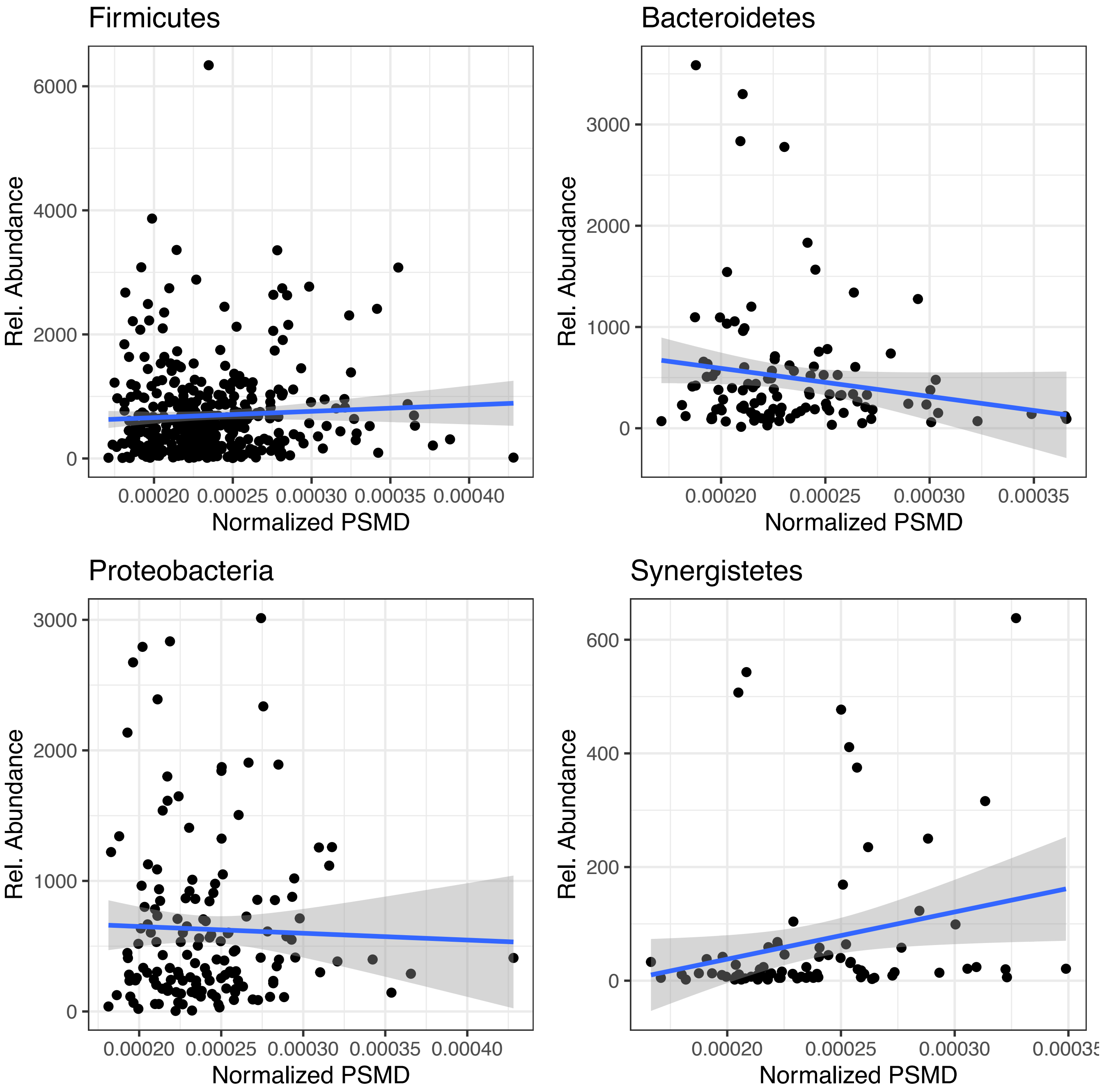

**Figure S6: Multivariate association analysis:** Scatter plots of PSMD depicting statistically significant associations at the phylum level

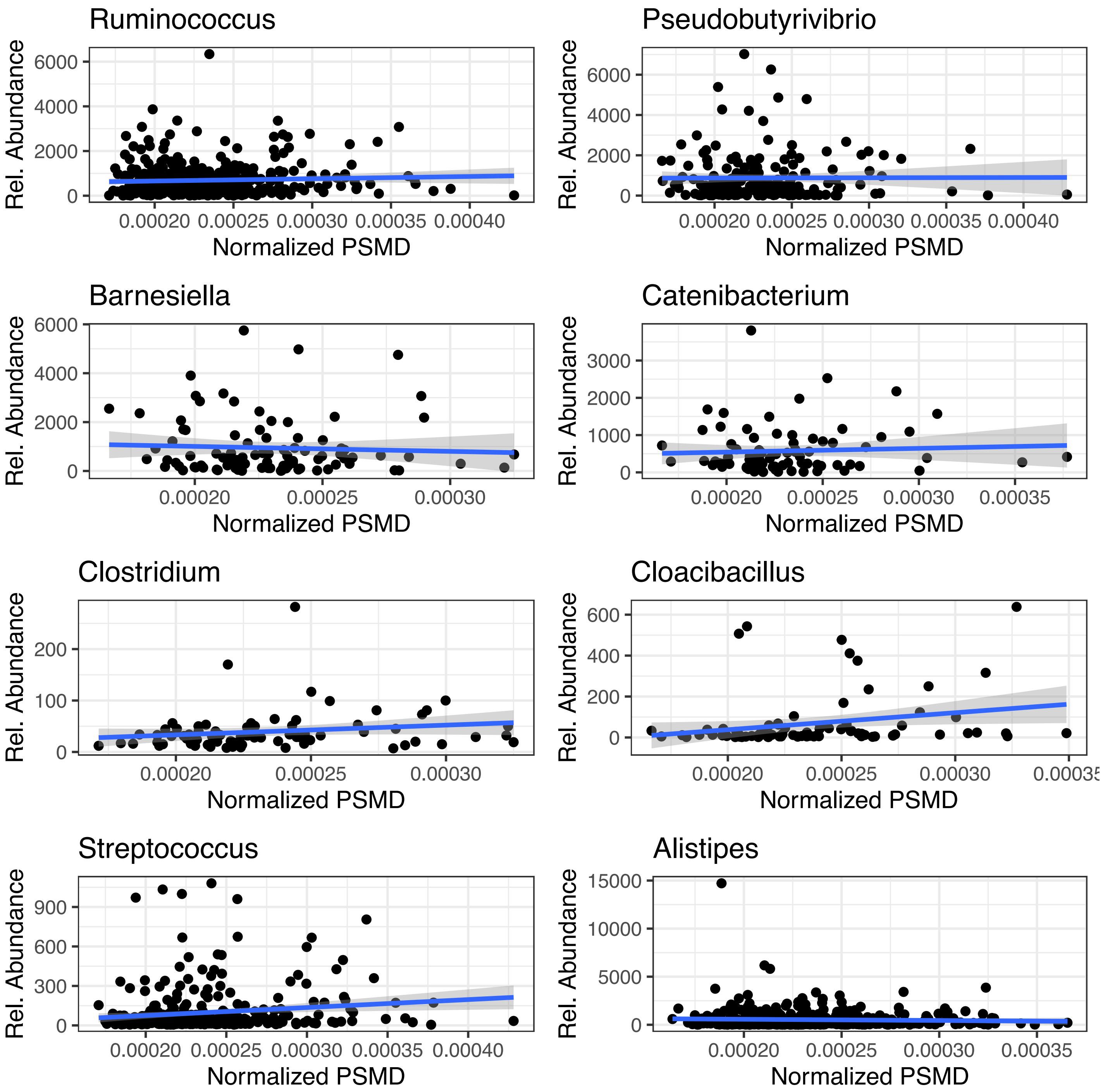

**Figure S7: Multivariate association analysis:** Scatter plots of PSMD depicting statistically significant associations at the genus level

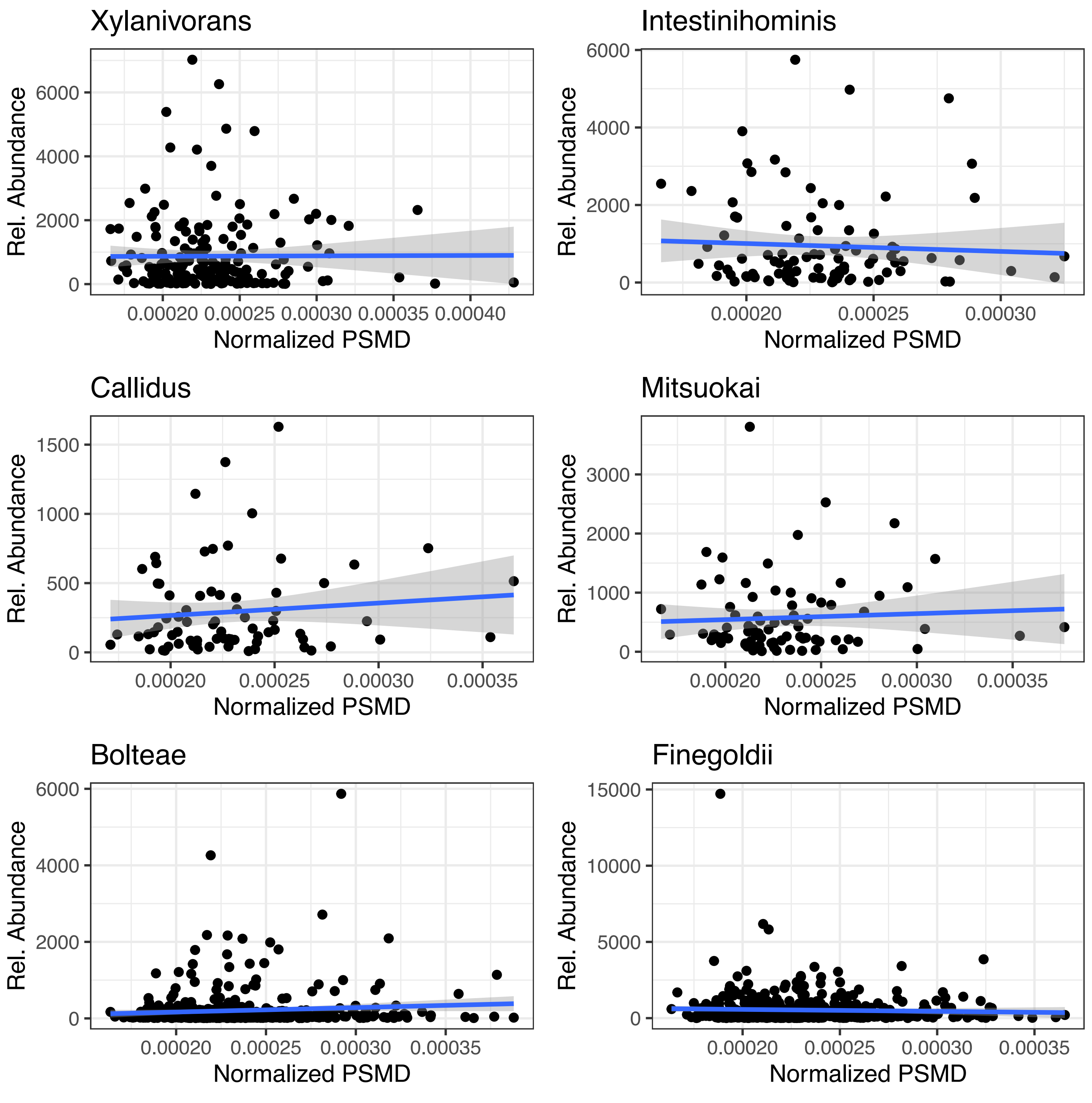

**Figure S8: Multivariate association analysis:** Scatter plots of PSMD depicting statistically significant associations at the species level

#### 2.2.2 Scatter plots of significant taxa associated with WMH

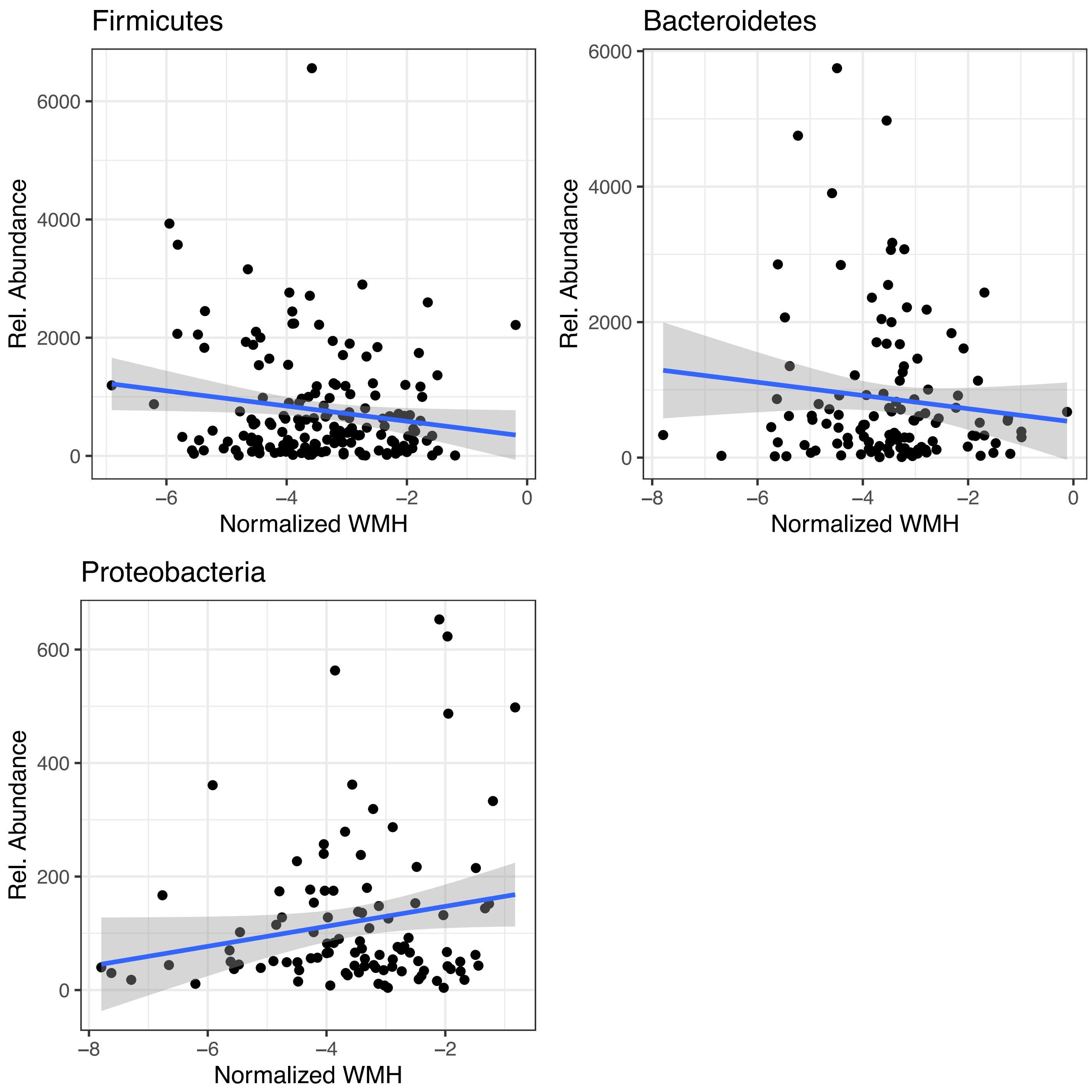

**Figure S9: Multivariate association analysis:** Scatter plots of WMH depicting statistically significant associations at the phylum level

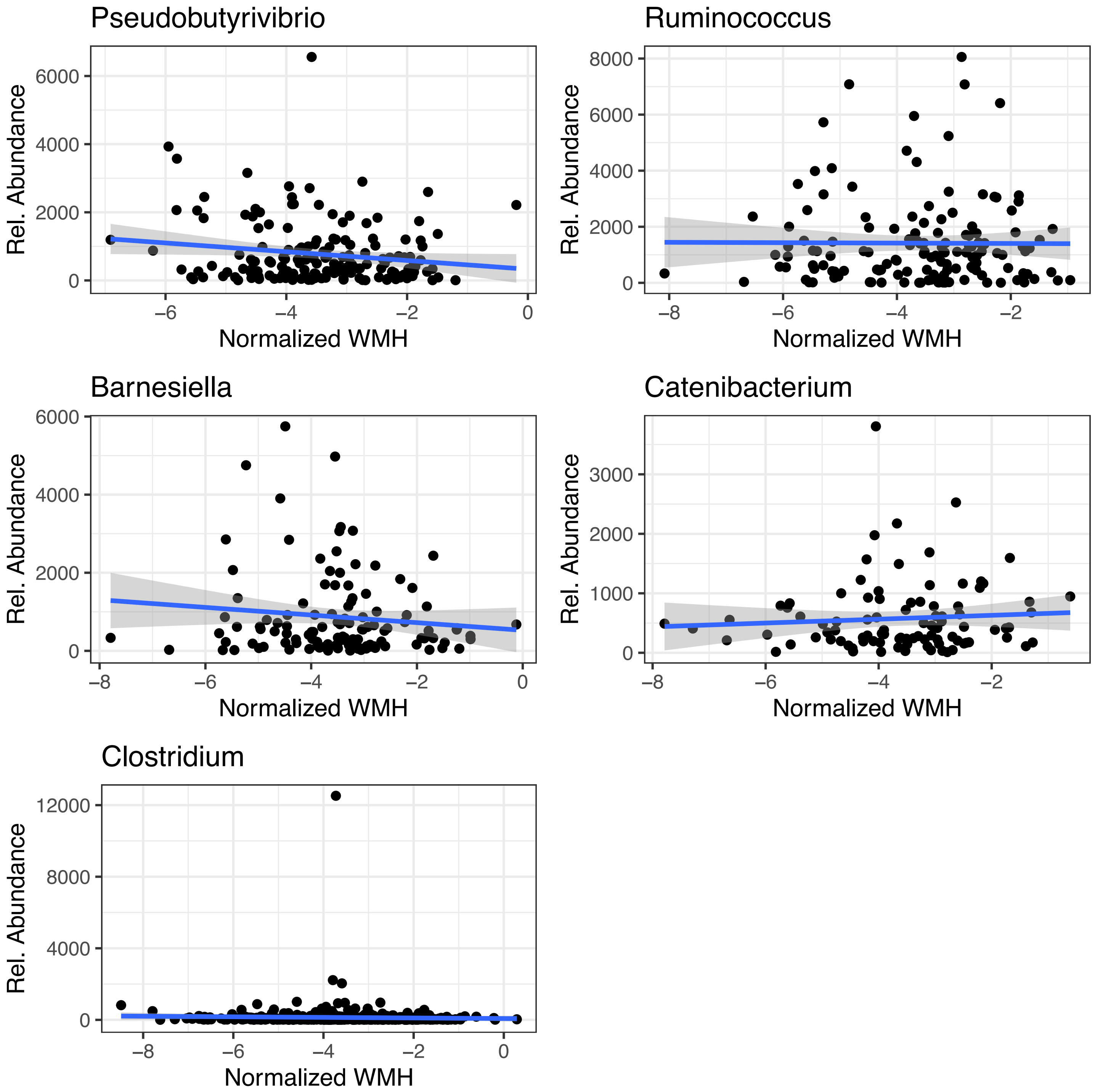

**Figure S10: Multivariate association analysis:** Scatter plots of WMH depicting statistically significant associations at the genus level

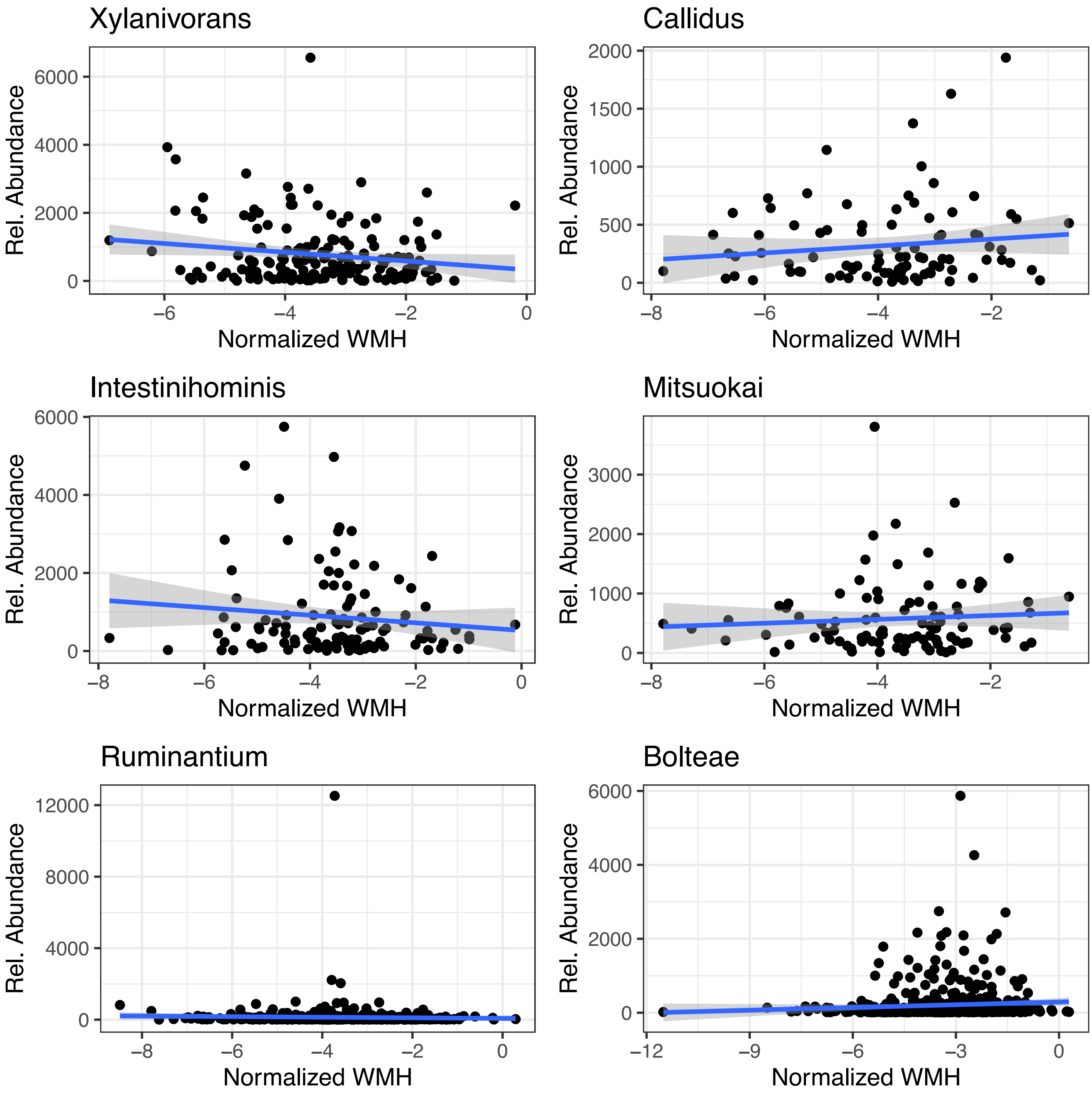

**Figure S11: Multivariate association analysis:** Scatter plots of WMH depicting statistically significant associations at the species level

##### 2.2.3 Scatter plots of significant taxa associated with EF

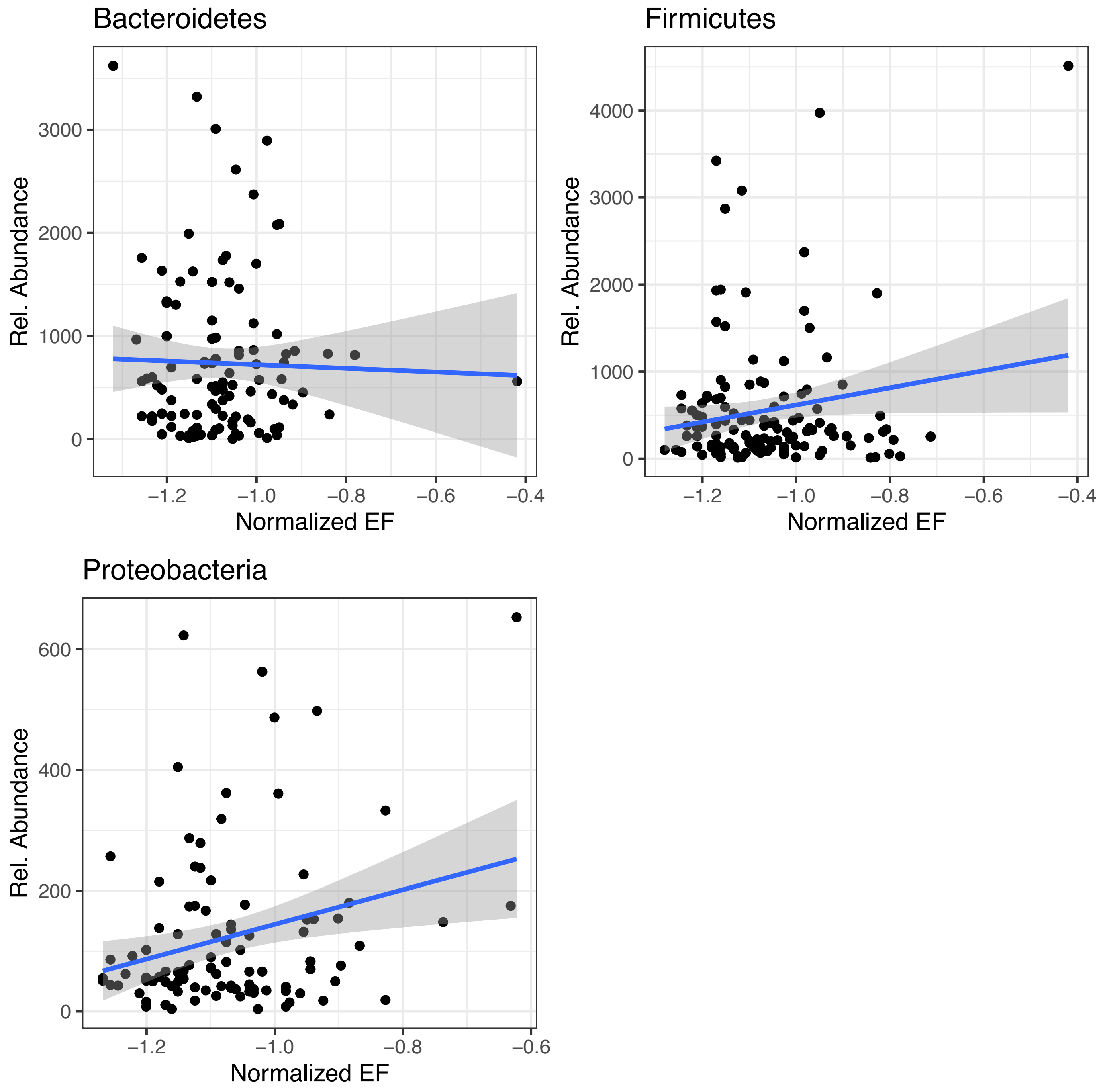

**Figure S12: Multivariate association analysis:** Scatter plots of EF depicting statistically significant associations at the phylum level

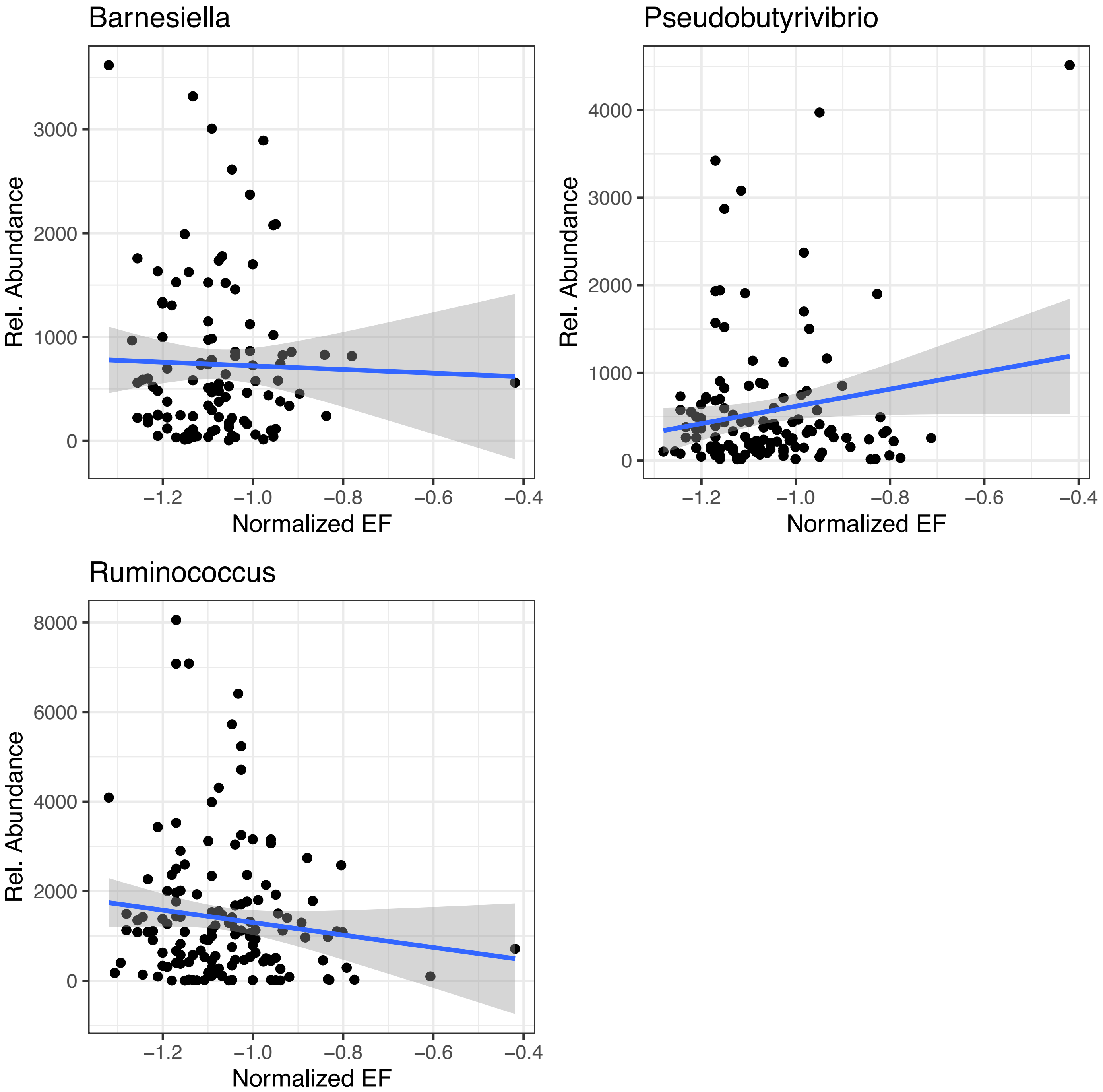

**Figure S13: Multivariate association analysis:** Scatter plots of EF depicting statistically significant associations at the genus level

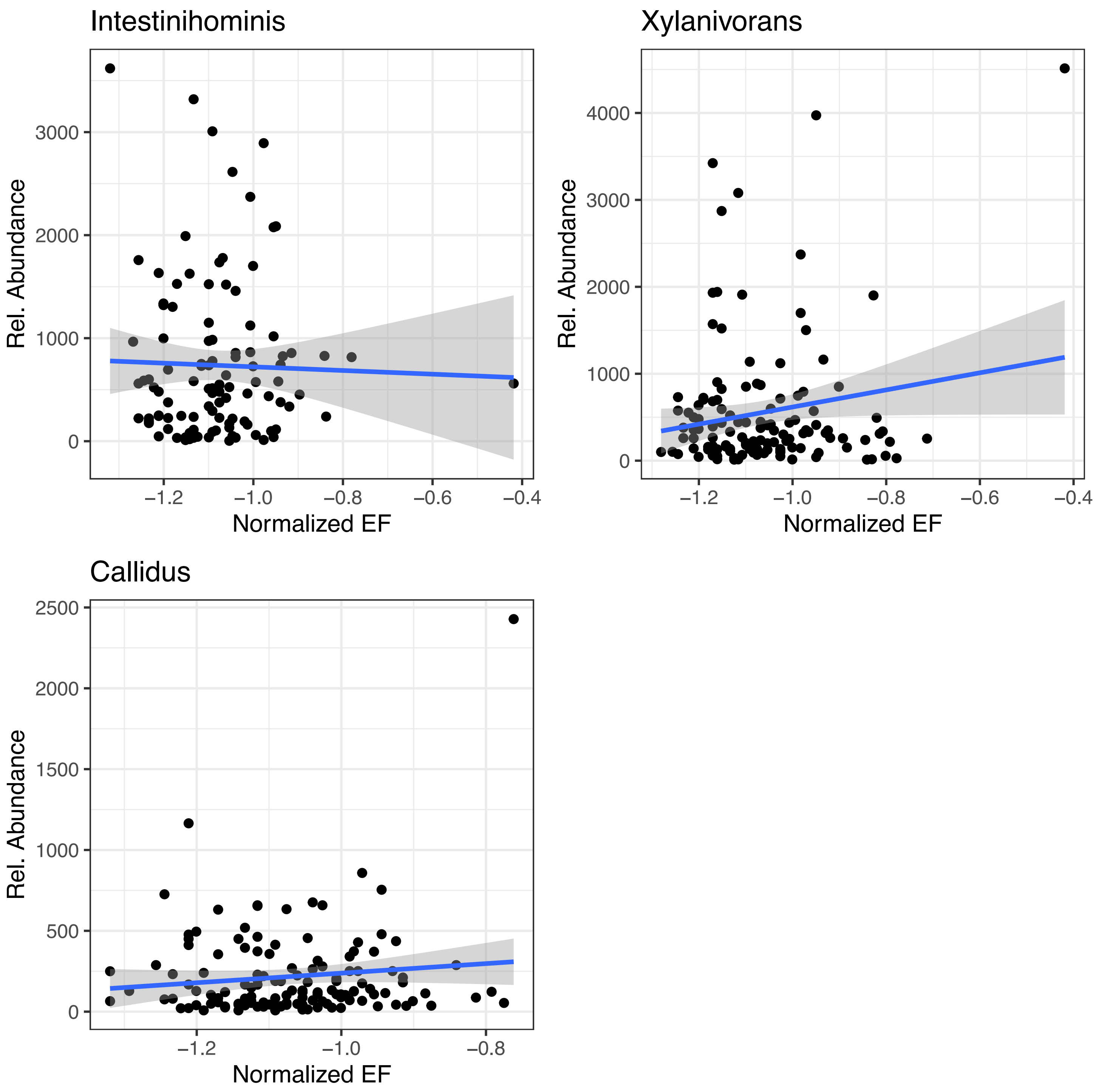

**Figure S14: Multivariate association analysis:** Scatter plots of EF depicting statistically significant associations at the species level

### 3. Differential abundance analysis

#### 3.1 Differentially abundant taxa, stratified by PSMD measures

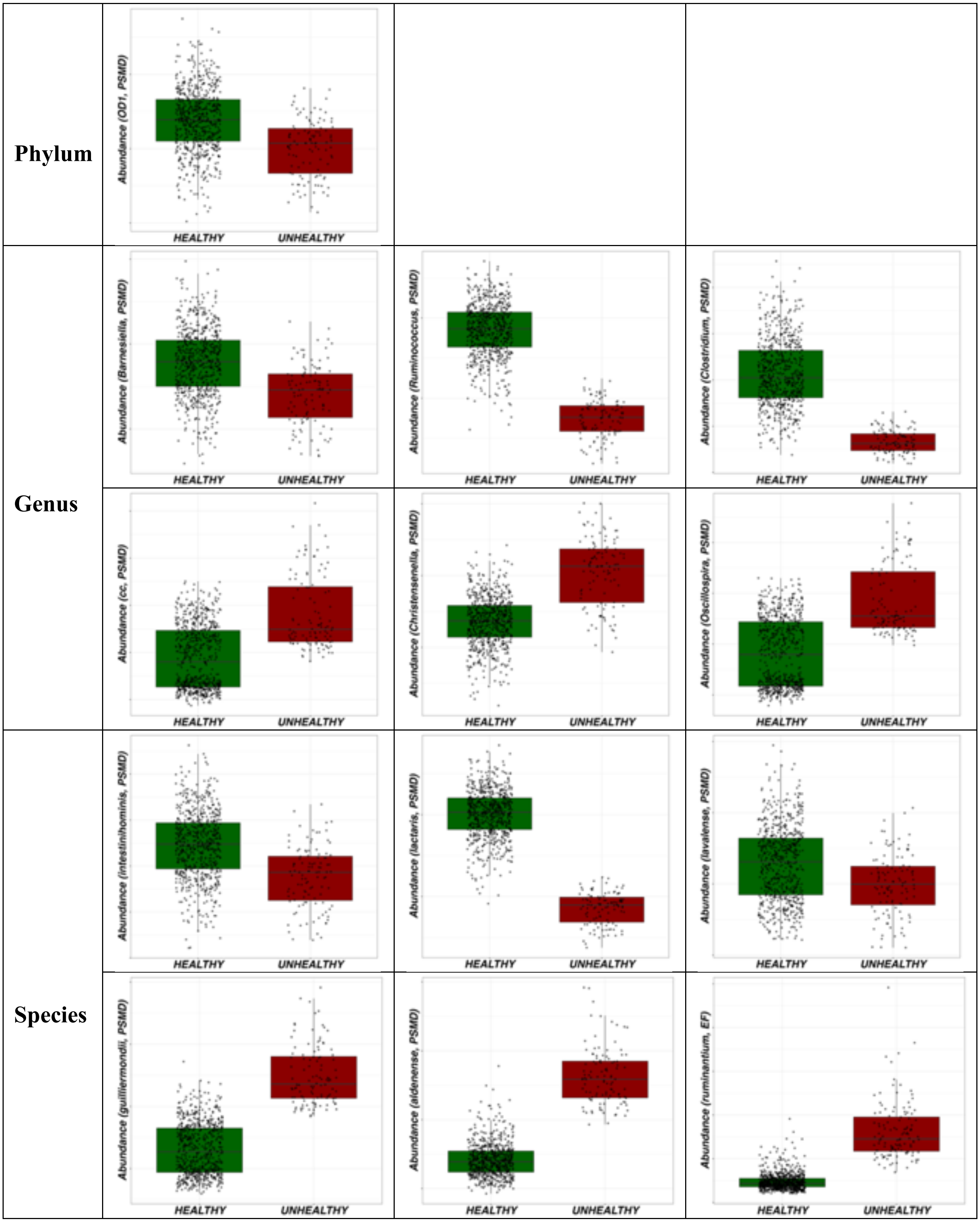

**Figure S15: Differential abundance analysis:** Differential abundance of the gut microbiome and measures of PSMD (Healthy = Lower burden, Unhealthy = High burden).

#### 3.2 Differentially abundant taxa, stratified by WHM measures

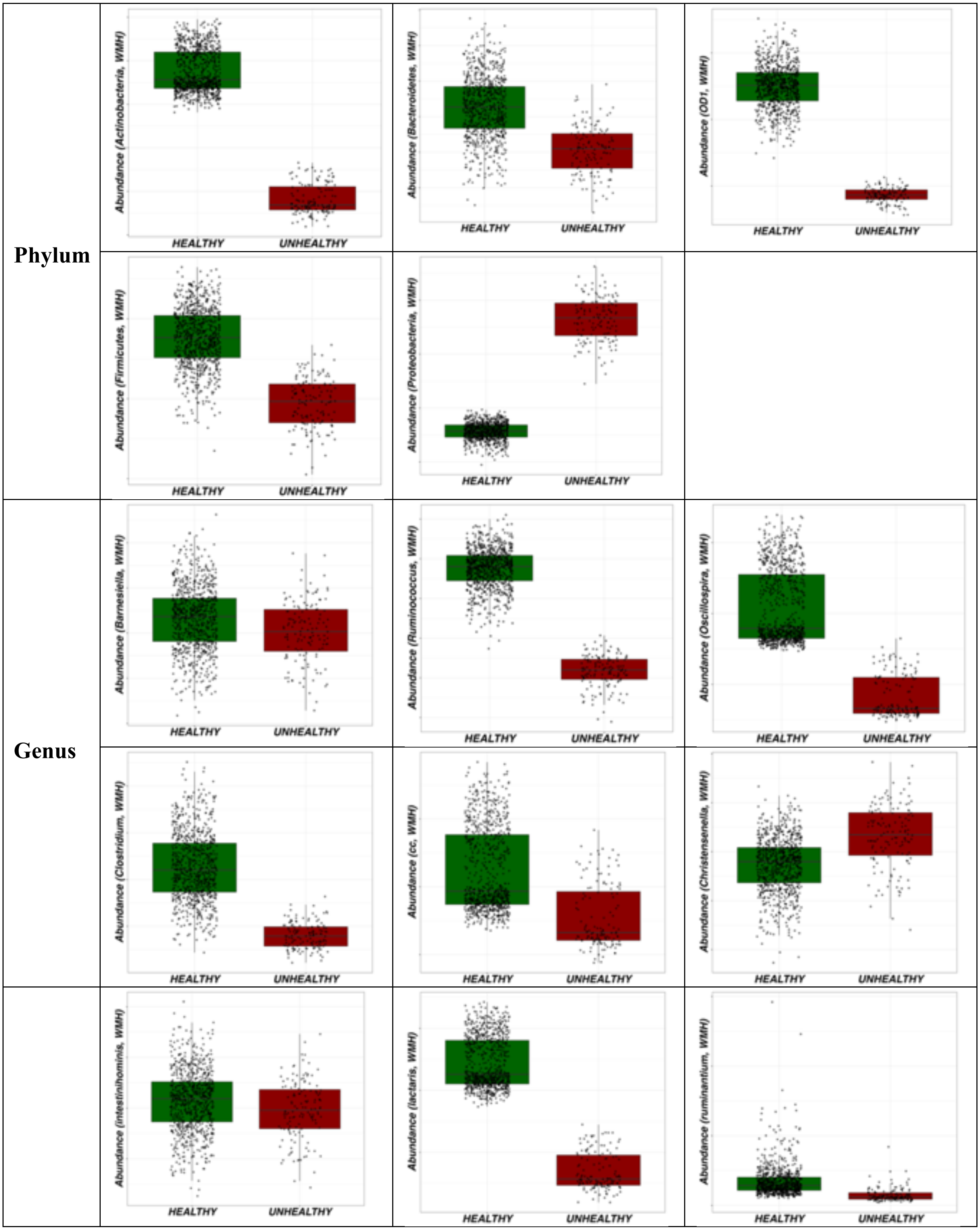
Phylum
Abundance (Actinobacteria, WMH)
HEALTHY
UNHEALTHY
Abundance (Bacteroidetes, WMH)
HEALTHY
UNHEALTHY
Abundance (OD1, WMH)
HEALTHY
UNHEALTHY
Abundance (Firmicutes, WMH)
HEALTHY
UNHEALTHY
Abundance (Proteobacteria, WMH)
HEALTHY
UNHEALTHY
Genus
Abundance (Barnesiella, WMH)
HEALTHY
UNHEALTHY
Abundance (Ruminococcus, WMH)
HEALTHY
UNHEALTHY
Abundance (Oscillospira, WMH)
HEALTHY
UNHEALTHY
Abundance (Clostridium, WMH)
HEALTHY
UNHEALTHY
Abundance (cc, WMH)
HEALTHY
UNHEALTHY
Abundance (Christensenella, WMH)
HEALTHY
UNHEALTHY
Abundance (intestinihominis, WMH)
HEALTHY
UNHEALTHY
Abundance (lactaris, WMH)
HEALTHY
UNHEALTHY
Abundance (ruminantium, WMH)
HEALTHY
UNHEALTHY

| | | |
|---|---|---|
| **Species** | 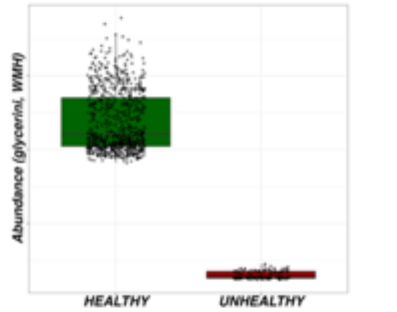 | 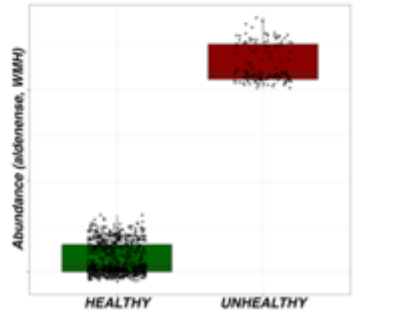 |

**Figure S16: Differential abundance analysis:** Differential abundance of the gut microbiome and measures of WMH (Healthy = Lower burden, Unhealthy = High burden).

#### 3.3 Differentially abundant taxa, stratified by EF measures

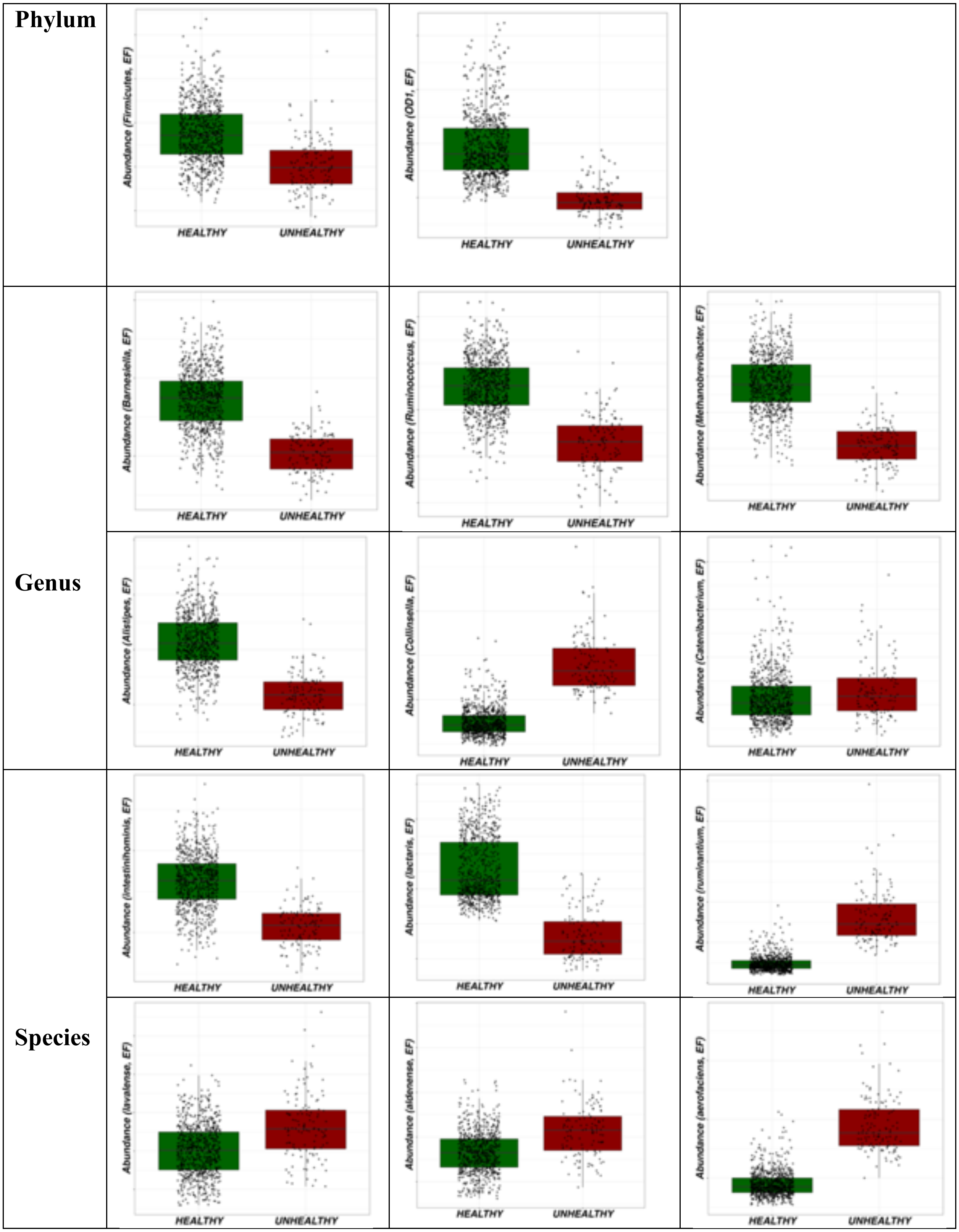

**Figure S17: Differential abundance analysis:** Differential abundance of the gut microbiome and measures of EF (Healthy = Lower burden, Unhealthy = High burden).

### 4. Diversity association analysis

#### 4.1 Alpha diversity

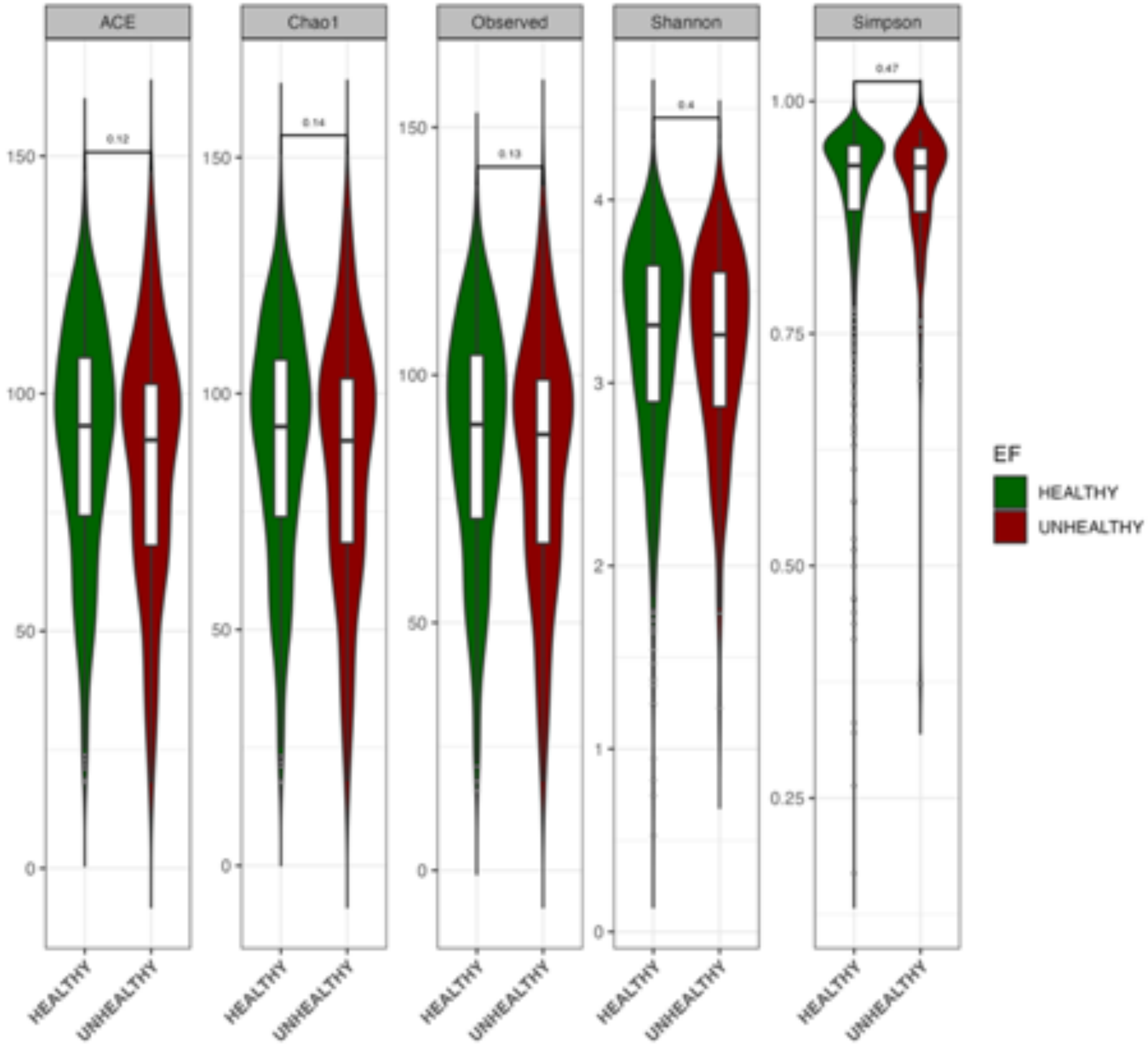

**Figure S18: Alpha diversity measures (ACE, Chao1, Observed, Shannon, and Simpson indexes) for EF stratified by burden groups.** No statistically significant differences were found for all measures, indicating a relatively stable gut microflora composition between the two groups. Significantly different between groups (Healthy = Lower burden, Unhealthy = High burden).

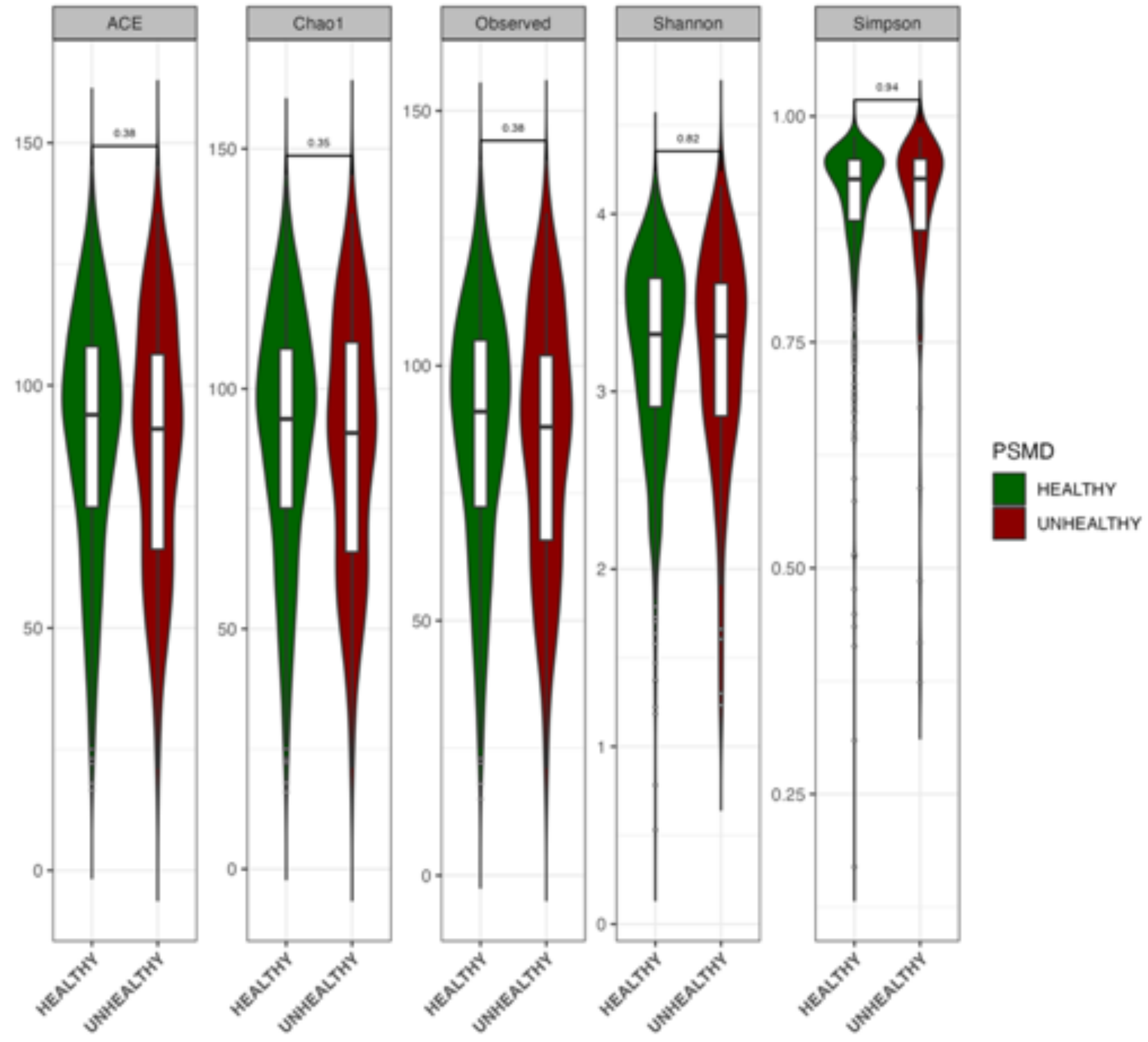

**Figure S19: Alpha diversity measures (ACE, Chao1, Observed, Shannon, and Simpson indexes) for PSMD stratified by burden groups.** No statistically significant differences were found for all measures, indicating a relatively stable gut microflora composition amongst the two groups (Healthy = Lower burden, Unhealthy = High burden).

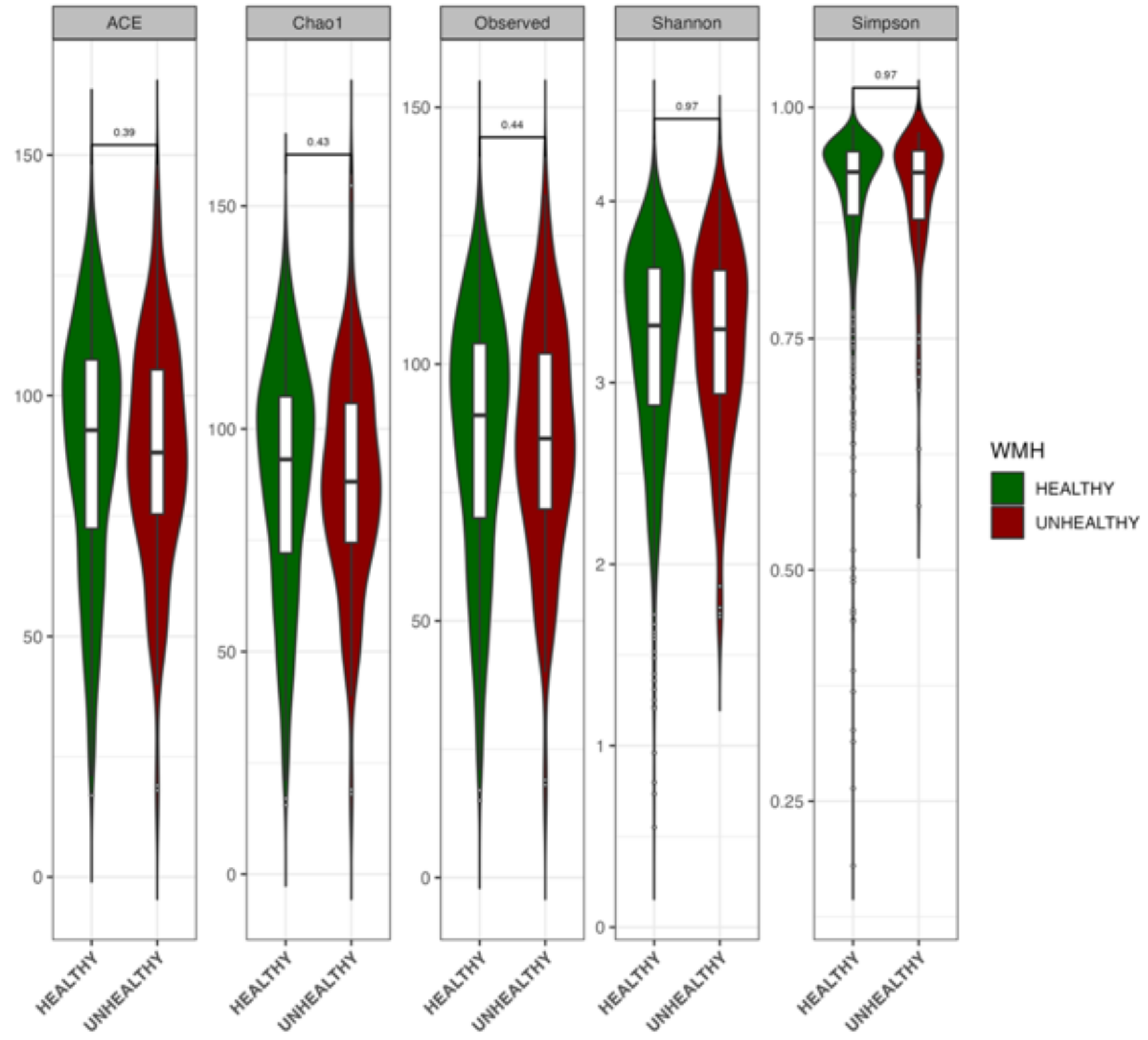

**Figure S20: Alpha diversity measures (ACE, Chao1, Observed, Shannon, and Simpson indexes) for WMH stratified by burden groups.** No statistically significant differences were found for all measures, indicating a relatively stable gut microflora composition amongst the two groups (Healthy = Lower burden, Unhealthy = High burden).

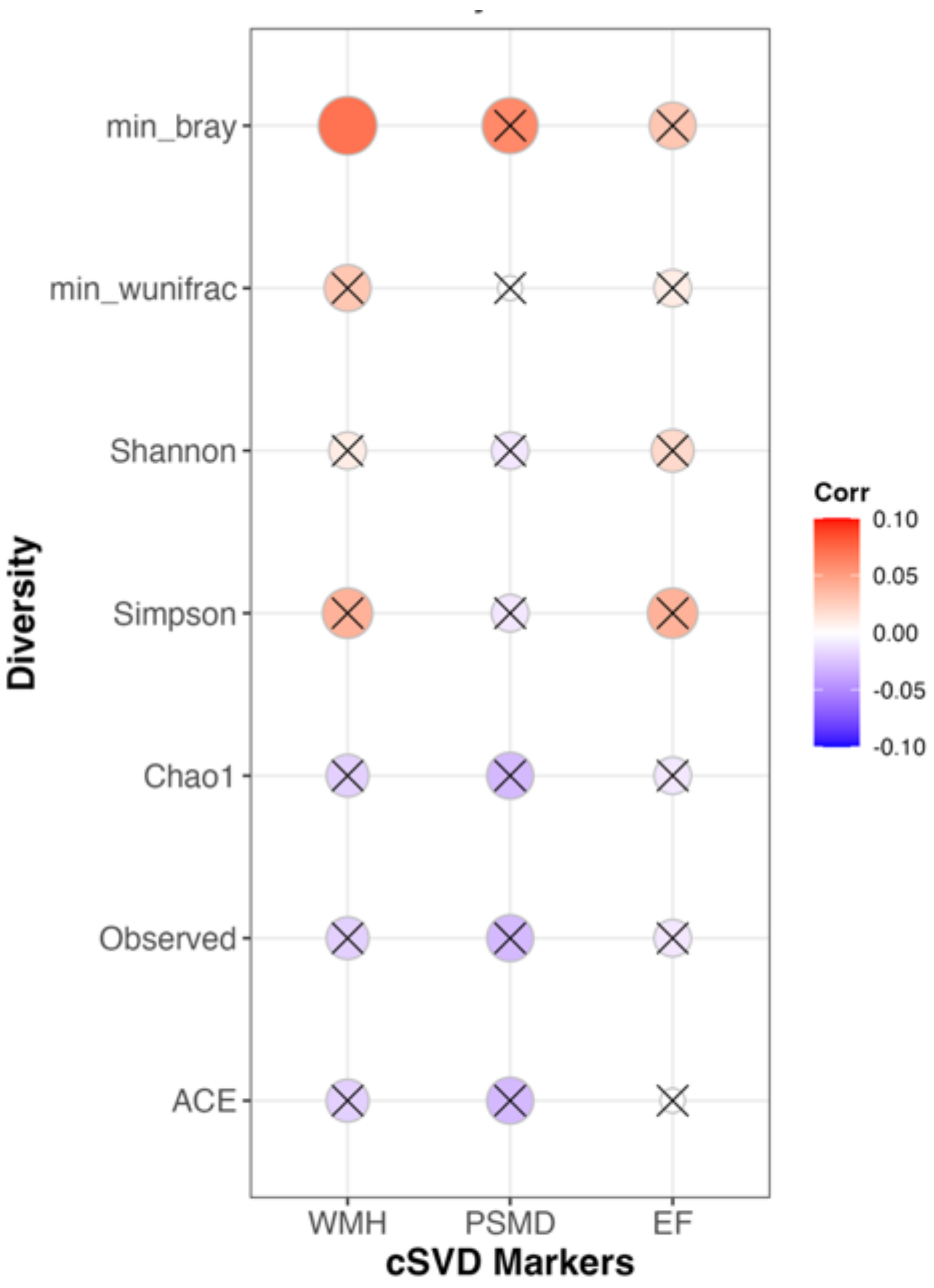

**Figure S21: Correlation between beta-diversity measures (min_bray, min_wunifrac), alpha-diversity (ACE, Observed, Chao1, Simpson, and Shannon indexes), and cSVD markers after adjusting for age, age2, sex, BMI, and the time difference between the stool collection and MRI scans**. No statistically significant correlations were found except for the association between WMH and minimum Bray-Curtis (min_bray). Just as in the stratified analysis, alpha diversity measures did not associate with markers of cSVD. Crosses in the plot indicate that a correlation was not significant (unadjusted p-value >0.05); min_bray and min_wunfrac are defined in the text.

#### 4.2 Beta diversity

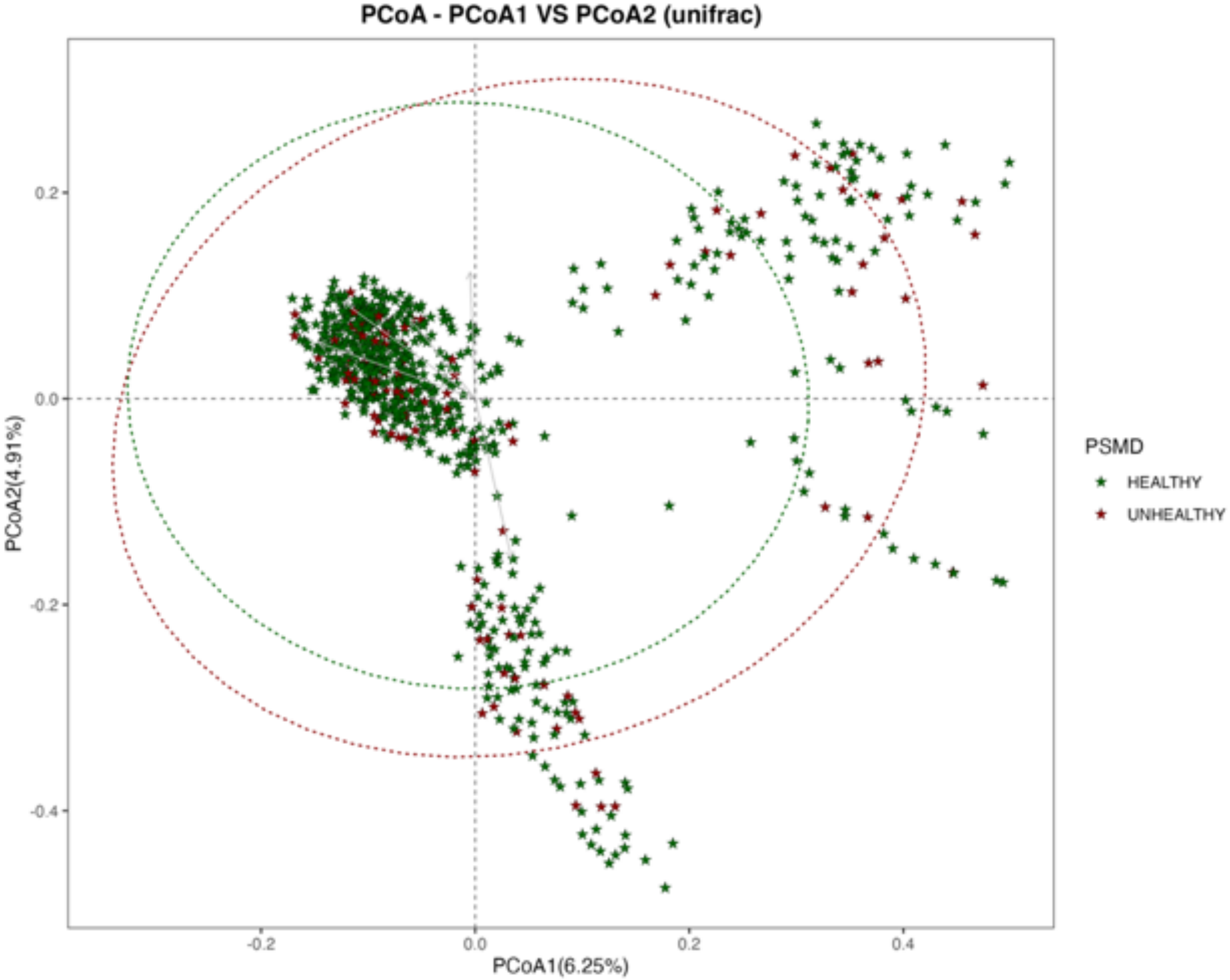

**Figure S22: Principal Coordinate Analysis (PCoA) depicting the diversity distribution between the different PSMD burden group**s. The different groups exhibit similar distribution. PCoA plot of unifrac distances for samples between WMH burden groups in microbial analysis (Healthy = Lower burden, Unhealthy = High burden).

### 5. Functional analysis with PICRUSt

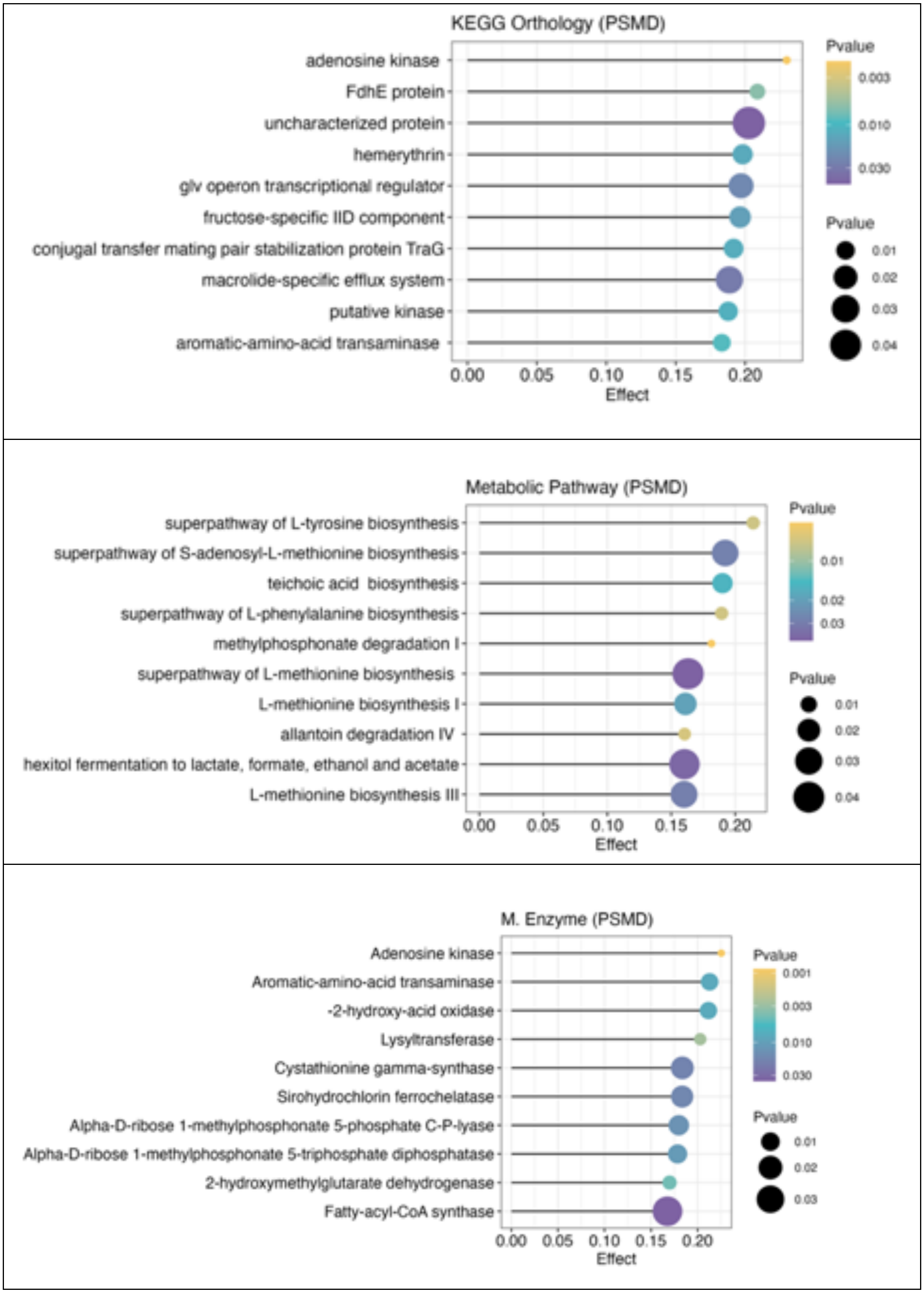

**Figure S23: Predicted functional role of the microbial communities associated with PSMD**

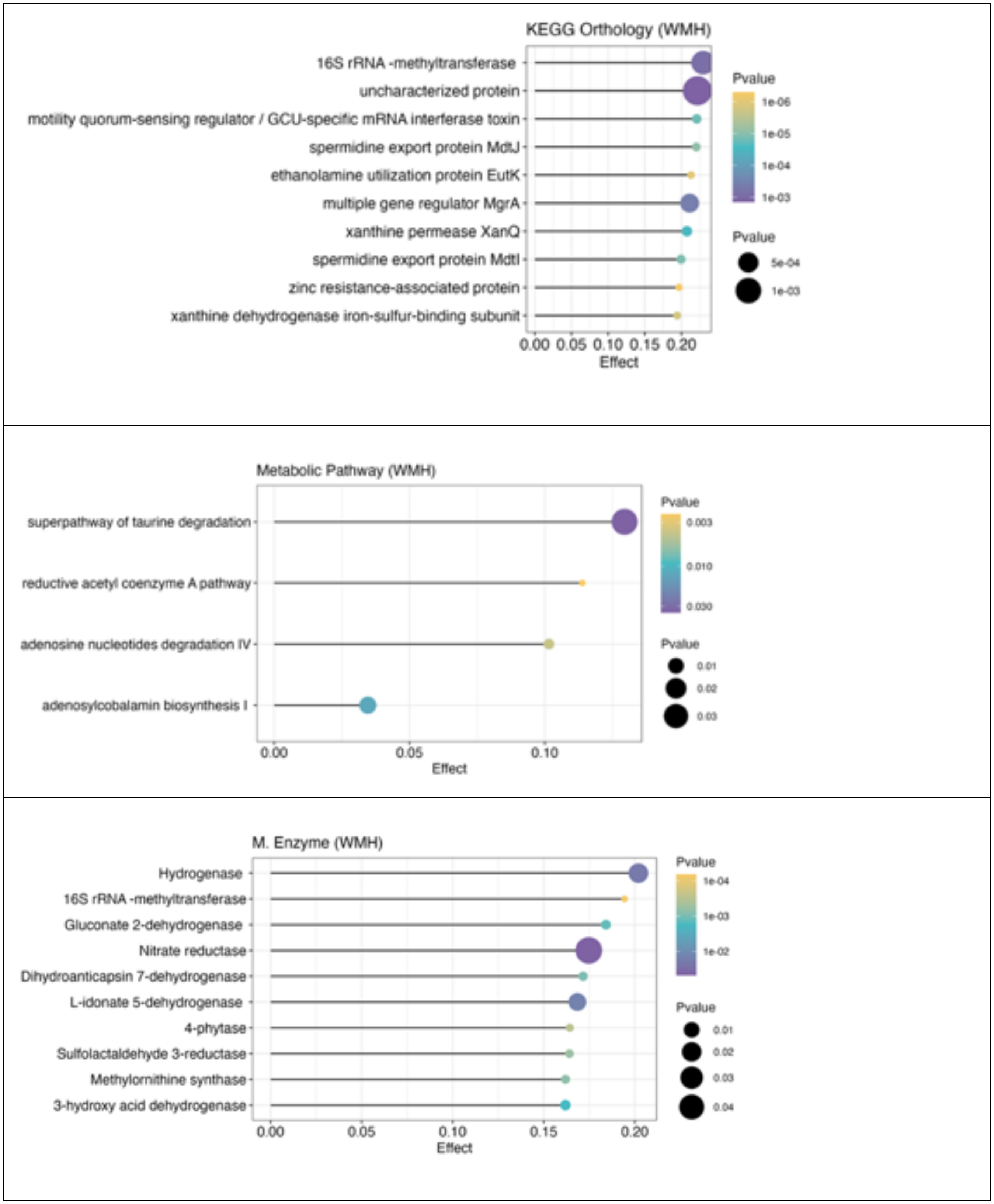

**Figure S24: Predicted functional role of the microbial communities associated with WMH**

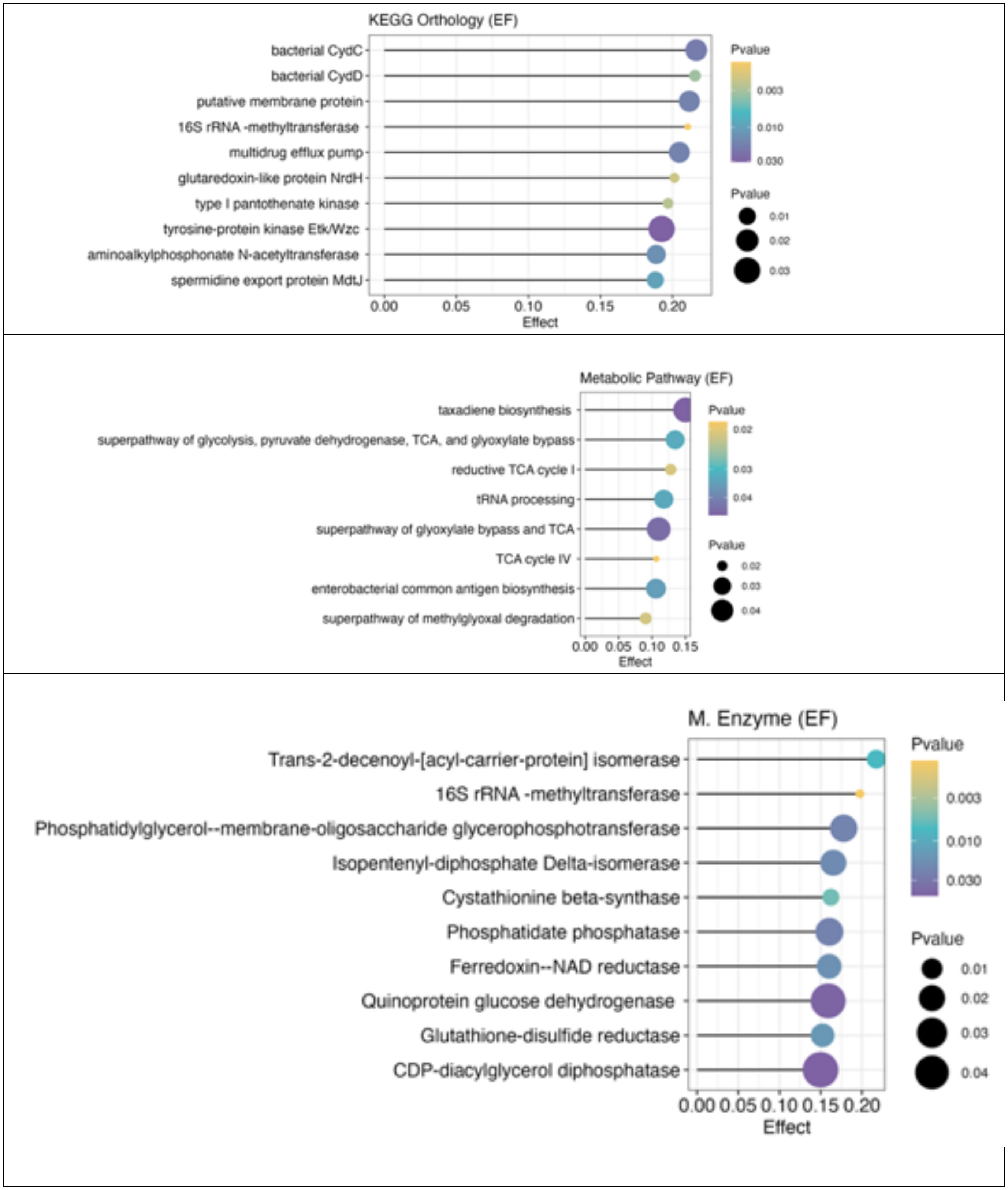

**Figure S25: Predicted functional role of the microbial communities associated with E**
